# The chromatin reader protein MLLT1 is critical to maintain normal B lymphopoiesis

**DOI:** 10.64898/2026.08.08.743534

**Authors:** Janani Prakash, Nicholas J. Achille, Emmalee R. Adelman, Shubin Zhang, John H. Bushweller, Maria E. Figueroa, Charles S. Hemenway, Nancy J. Zeleznik-Le

## Abstract

MLLT1 (also named ENL) is a chromatin reader protein whose encoding gene was originally identified as a chromosomal translocation partner with *MLL*(*KMT2A*) in acute leukemia. However, its role in normal hematopoiesis has not been investigated. This study uncovers a critical role of Mllt1 in normal B cell lymphopoiesis. We found Mllt1 to be essential for early B lymphocyte development using a conditional *Mllt1* knockout mouse model that we developed. A significant decrease of bone marrow B-lineage progenitors, splenic transitional B cells and peripheral blood B cells were observed in *Mllt1^del^* mice compared to control *Mllt1^fl/fl^* mice. Similarly, *Mllt1* deletion in *in vitro* cultured B-enriched progenitor cells from *Mllt1^fl/fl^; Rosa26CreER^T2/+^* mice resulted in reduced B cells, demonstrating the cell-intrinsic role of *Mllt1* in this process. Direct MLLT1 target genes including *Il7r* and critical B-lineage transcription factors, *Ebf1* and *Pax5*, were decreased following *Mllt1* deletion. Gene set enrichment, gene ontology, and functional analyses of *Mllt1*-deficient cells showed significant alterations related to B cell development, critical relevant signaling pathways, DNA replication, and mitochondrial function. *In vitro* complementation with *MLLT1* rescued the B cell phenotype observed with endogenous *Mllt1* deletion; however, specific MLLT1 YEATS domain mutants lacking chromatin reader and RNA-binding functions were unable to rescue the phenotype. Taken together, our research demonstrates a previously unappreciated role for MLLT1 as critical for maintenance of B cell lymphopoiesis.

## Introduction

Hematopoiesis, the process by which all types of blood cells are formed, is complex and highly regulated. Epigenetic mechanisms play a significant role in the initiation or silencing of gene expression pathways to guide hematopoietic stem cells (HSCs) to develop into specific blood lineages. Disruption in the function of chromatin readers, writers and erasers can lead to a defective chromatin landscape and abnormal hematopoiesis. The YEATS (Yaf9, ENL, AF9, Taf14, Sas5) domain-containing proteins are one group of chromatin reader proteins that recognize and bind to specifically acylated histone residues through their conserved N-terminal YEATS domain [1, 2]. Two of these proteins were originally identified as fusion partners of MLL/KMT2A: ENL/MLLT1 and AF9/MLLT3. The MLL-ENL fusion oncoprotein gives rise to predominantly B-lineage or mixed phenotype acute leukemia, while MLL-AF9 is more commonly associated with acute myeloid leukemia [3]. These fusion oncoproteins and the wildtype ENL and AF9 proteins have been primarily studied for their roles in leukemia [4, 5].

ENL/MLLT1 and AF9/MLLT3 are very close paralogs with high sequence conservation of their functional domains. Both MLLT1 and MLLT3 recognize and bind to crotonylated or acetylated histone H3 lysine residues and also bind RNA through their N-terminal YEATS domains [1, 2, 6]. MLLT1 and MLLT3 both regulate transcription of direct target genes by recruiting several different transcription regulatory complexes which compete for binding to their C-terminal domain [7]. These include the Super Elongation Complex (SEC), the Dot1L H3K79 methyltransferase complex, and the Polycomb Repressive Complex 1 (PRC1) [8–11]. Even though MLLT1 and MLLT3 have similar functional domains they are differentially expressed in specific hematopoietic cell subsets. MLLT3 is very abundant in long-term hematopoietic stem cells and has been identified as critical to maintain functional LT-HSCs [12]. In contrast, MLLT1 is not expressed at a high level in LT-HSCs, but rather expressed in different progenitors, immature and mature hematopoietic subsets [13]. Among the hematopoietic cells that express *Mllt1* in the bone marrow, subpopulations of B-lineage cells exhibit high expression levels suggesting the possibility of regulatory function by Mllt1 in developing B cells. Apart from its role in leukemia, the function of Mllt1 in normal hematopoiesis has not been investigated. In this study, we identified a novel function of Mllt1 as a positive regulator of early B-cell development. Mechanistically, it regulates the expression of critical transcription factors and components of signaling pathways. It also supports cellular proliferation and mitochondrial function for normal B cell development.

## Methods

### Mice

*Mllt1* conditional knock-out mice (*Mllt1^fl/fl^; Rosa26Cre-ER^T2/+^* mice (experimental) or *Mllt1^fl/fl^; +/+* (control)) were generated that result in cre-mediated excision of the third exon of *Mllt1* and absence of detectable *Mllt1* transcripts following administration of tamoxifen to adult animals or exposure to 4-hydroxy-tamoxifen (4OHT) to cells *in vitro*. Genotyping primers and more details are provided in Online Supplementary materials. All animal studies were performed according to standards set forth in the NIH guidelines and approved by Loyola University’s Institutional Animal Care and Use Committee.

### Flow cytometry

Cells isolated from mouse tissues were incubated with 1X RBC lysis buffer. Cells from both *in vivo* and *in vitro* experiments were stained with Live/Dead Zombie Violet dye (Biolegend, cat# 423114), washed with FACS buffer (PBS+2%FBS) and incubated with fluorescent conjugated-lineage-specific antibodies (*Online Supplementary Table S2*) in brilliant stain buffer (BD Horizon, cat# 563794) or FACS buffer. As appropriate, cells were stained for Mitotracker and/or proliferation marker. Cells were analyzed by flow cytometry using LSR Fortessa cytometer (Becton Dickinson) followed by analysis using FlowJo v9 and v10 software. Cell sorting was performed by flow cytometry core staff using Cytek Aurora CS.

### In vitro progenitor B cell differentiation assay

B-enriched lineage negative cells were isolated from bone marrow cells from *Mllt1^fl/fl^; Rosa26Cre-ER^T2/+^* or *Mllt1^fl/+^; Rosa26Cre-ER^T2/+^* mice by negative selection using MagniSort mouse B cell enrichment kit (ThermoFisher scientific, cat# 8804-6827-74) as per manufacturer’s protocol. Resulting B-enriched cells were equally plated between control (400nM ethyl alcohol) and knock-out (400nM 4OHT) for culture.

### RNA sequencing

For cells cultured *in vitro*, total RNA was extracted from each of vehicle- and 4OHT- treated B-enriched lineage negative bone marrow cells of three independent biological samples using TRI Reagent (Sigma-Aldrich, #T9424), according to the manufacturer’s instructions. Similarly for *in vivo* samples, RNA was isolated from FACS isolated mouse bone marrow-derived B progenitors and CLP from each of three tamoxifen treated *Mllt1^fl/fl^; Rosa26Cre-ER^T2/+^* and *Mllt1^fl/fl^* mice. RNA sequencing for *in vitro* samples was performed by Novogene (Sacramento, CA), and *in vivo* samples were prepared by the University of Miami Hussman Institute for Human Genomics Sequencing Core. Detailed methods are provided in *Online Supplemental Methods*.

### Quantitative RT-PCR

Methods are provided in Supplement.

### Seahorse Assay

Cellular oxygen consumption rate (OCR) was measured using the Seahorse XFe96 Analyzer (Agilent) for lineage negative cells isolated from the bone marrow of *Mllt1^del^* and control mice.

Molecular cloning of MLLT1 and its mutants and *in vitro* complementation assay described in Supplemental methods.

### Statistical analysis

The statistical analyses of all data were calculated using GraphPad Prism v10. Comparisons between two groups were performed using One sample t test when control is normalized to 100 or 1, otherwise done by an unpaired, two-tailed Student’s t-test. Data are presented as mean ± standard deviation (SD), with statistically significant p-values are represented as \**p* < 0.05, \*\**p* < 0.01, \*\*\**p* < 0.001, \*\*\*\**p* < 0.0001. Sample sizes (n) and any statistical test which was applied other than the listed methods are specified in the figure legends.

## Results

### Mllt1 is required to maintain B cells in vivo

While *Mllt1* is expressed in multiple hematopoietic lineages, it is expressed relatively higher in subpopulations of B-lineage cells suggesting the possibility of regulatory functions by Mllt1 in developing bone marrow B cells (GSE15907, Immunological Genome Project gene expression database) [13]. Given that constitutive *Mllt1* deletion caused early mortality in mouse embryos precluding detailed analysis [14], we generated mice with a conditional deletion *Mllt1^fl/fl^; Rosa26CreER^T2/+^* or control *Mllt1^fl/fl^* to study the role of *Mllt1* in B cell development *in vivo* following tamoxifen injection.

Immunophenotyping of bone marrow hematopoietic cells at 2 weeks post *Mllt1* deletion demonstrated a significant decrease of total B cells (Figure 1A; *Online Supplementary Figure S1A*). Furthermore, comprehensive analysis of different stages of developing B cells in the bone marrow revealed that a differentiation block occurs at the Pre Pro-B to Pro-B transition with a pronounced reduction of early B-lineage progenitors including Pro-B, Early Pre-B, Late Pre-B, lmmature B, and Transitional B populations (Figures 1A- D). In contrast, other bone marrow cell populations such as stem cells (LT-HSC, ST-HSC), multipotent progenitors (MPPs) and myeloid populations (CMP, GMP and MEP) were not consistently affected following *Mllt1* deletion (*Online Supplementary Figure S1D*). Similarly, there were no significant differences in other lymphocyte populations such as T cells and NK cells in *Mllt1^del^* mice (*Online Supplementary Figure S1D*). As the most significant and consistent phenotype was observed in B-lineage populations, we further investigated the role of *Mllt1* in the B-lineage.

**Figure 1.**
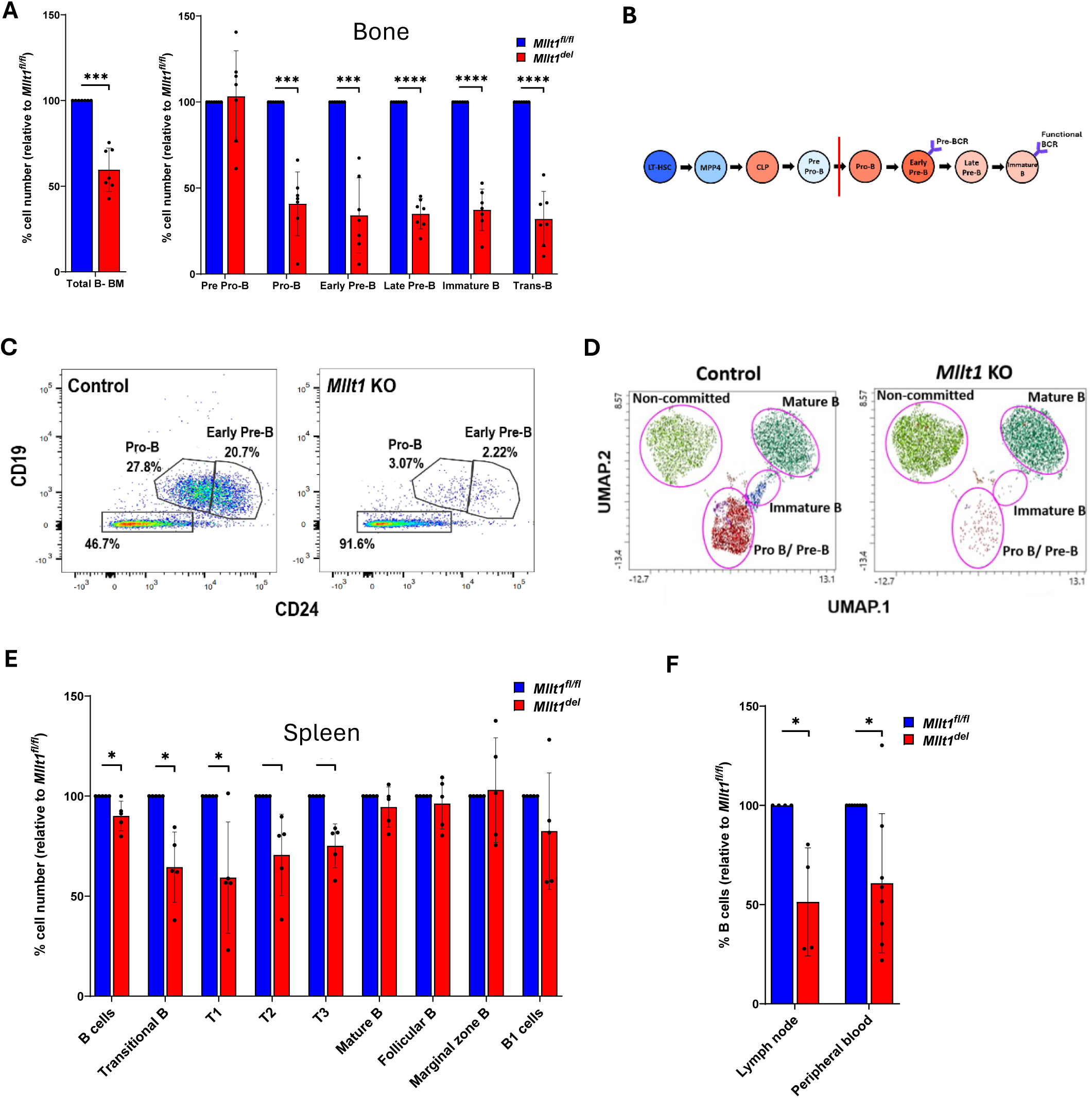
Loss of *Mllt1* Decreases the Number of B Cells *in vivo*. **(A)** Relative number of bone marrow B-lineage cell subpopulations following *in vivo Mllt1* deletion (n = 7). **(B)** Schematic representation of bone marrow B-cell development with color indicating relative *Mllt1* expression levels (blue= low; red=high). Red line highlights the stage at which *Mllt1* deletion has the most dramatic effect. **(C)** Representative flow plot of CD19 and CD24 surface expression in *Mllt1^fl/fl^* control and *Mllt1^del^* bone marrow cells to identify Pro-B and Early Pre-B from B220+ CD43+ gated population as determined by flow cytometry. **(D)** UMAP plot shows different populations of committed and non-committed B cells between *Mllt1^fl/fl^* control and *Mllt1^del^* (KO) mouse. **(E)** Relative number of spleen B-cell subpopulations following *in vivo Mllt1* deletion (n = 5). **(F)** Relative number of recirculating B cells of inguinal lymph node (n = 4) and peripheral blood (n = 6) following *in vivo Mllt1* deletion. Data shown as mean ± SD. *p < 0.05, **p < 0.01, ***p < 0.001, ****p < 0.0001 by One sample t test.

To evaluate the fate of transitional B cells exiting bone marrow to the spleen for further maturation, splenic B cells were immunophenotyped. Splenic total B and transitional B cells were markedly reduced 2 weeks post *Mllt1* deletion (Figure 1E). However mature B cells including follicular and marginal zone B cells, and B1 cells were not significantly altered (Figure 1E). In addition, spleen weight to body weight ratio was not affected by *Mllt1* deficiency (*Online Supplementary Figure S1B*). To investigate the effect of *Mllt1* loss on recirculating B cells, peripheral blood and inguinal lymph nodes were analyzed based on the surface expression of B220^+^ and CD19^+^. B cells in both peripheral blood and lymph nodes were noticeably decreased following loss of *Mllt1* (Figure 1F). Analysis of later time points following *Mllt1* deletion, specifically at 5 weeks, demonstrated that all bone marrow B-lineage progenitors and total B cells were recovered (*Online Supplementary Figure S1E*). Similar recovery of B cells was observed in spleen and lymph nodes by 5 weeks post *Mllt1* deletion (*Online Supplementary Figure S1E*). In contrast to the recovery of B cells observed in primary and secondary lymphoid organs, the peripheral blood still maintained B-lymphopenia 5 weeks post tamoxifen administration. This observation suggests that the phenotype observed in blood does not directly mirror the bone marrow phenotype (*Online Supplementary Figure S1E*).

When the developing B cells in the bone marrow were recovered at 5 weeks following *Mllt1* deletion, the number of B cells in the peripheral blood remained low relative to control. To comprehend the reason behind the persistence of B-lymphopenia as a separate phenotype from that in the bone marrow, we investigated the effect of *Mllt1* loss on the expression level of genes that encode receptors critical for the survival of peripheral blood B cells or the egress of B cells from the bone marrow/spleen. Baffr is the primary receptor for BAFF-mediated mature B cell survival and is necessary to maintain the peripheral blood pool [15, 16]. The expression of *Baffr* was significantly decreased in the bone marrow and peripheral blood of *Mllt1*^del^ mice (*Online Supplementary Figure S3A*). *S1pr1* encodes the receptor present on the surface of B cells and other mature cells to sense the sphingolipid S1P concentration in the circulation as an exit signal to regulate the egress of immature B cells from bone marrow and mature B cells from spleen into the circulation [17, 18]. We found that *S1pr1* expression trended lower in the bone marrow but not in the spleen of *Mllt1*^del^ mice compared to control (*Online Supplementary Figure S3B*). Because B cells decreased dramatically in the bone marrow and peripheral blood of *Mllt1*-deficient mice, plasma immunoglobulin levels were analyzed, and no significant changes were found in plasma immunoglobulin levels except for IgA which was decreased at 2 weeks post *Mllt1* deletion (*Online Supplementary Figure S1C*).

### In vitro model system demonstrates cell-intrinsic Mllt1 requirement for B lineage cell differentiation

*In vivo*, we found that *Mllt1* was required to maintain B cells starting at the Pro-B cell stage. To complement the *in vivo* system and to determine the cell-intrinsic role of Mllt1, we developed an *in vitro* B-cell differentiation assay where the end point of differentiation is at the Pro-B cell stage. When B-enriched lineage negative bone marrow cells were cultured with IL-7, Flt3-L and SCF, *Mllt1^del^* cells exhibited a robust phenotype of ∼70% - ∼80% fewer B, Pro-B and Pre Pro-B cells compared to vehicle (Figures 2A-D; *Online Supplementary Figure S2A*). To address any potential effect of 4- hydroxy tamoxifen, we performed the same experiment using *Mllt1^fl/fl^* cells and determined that it had no effect on B cell differentiation in this system (*Online Supplementary Figure S2E*). To address whether loss of one *Mllt1* allele could impact the differentiation of B-lineage cells, *Mllt1^fl/+^; Rosa26CreER^T2/+^* cells were tested in the same system. One allele deletion of *Mllt1 in vitro* gave rise to significantly fewer B cells than control; however, the extent of the effect was less than with loss of both *Mllt1* alleles (*Online Supplementary Figure S2F*). Interestingly, the B cell deficit favored the relative increase of myeloid cells (Monocytes: CD115+ and CD11b+; Granulocytes: Gr- 1+), not from their increased proliferation, but rather due to decreased proliferation of *Mllt1^del^* B cells (Figures 2C-D; *Online Supplementary Figure S2B-S2D*). Therefore, in the presence of the same external stimuli, the ability of cells to differentiate along the B cell lineage is impaired following *Mllt1* deletion.

**Figure 2.**
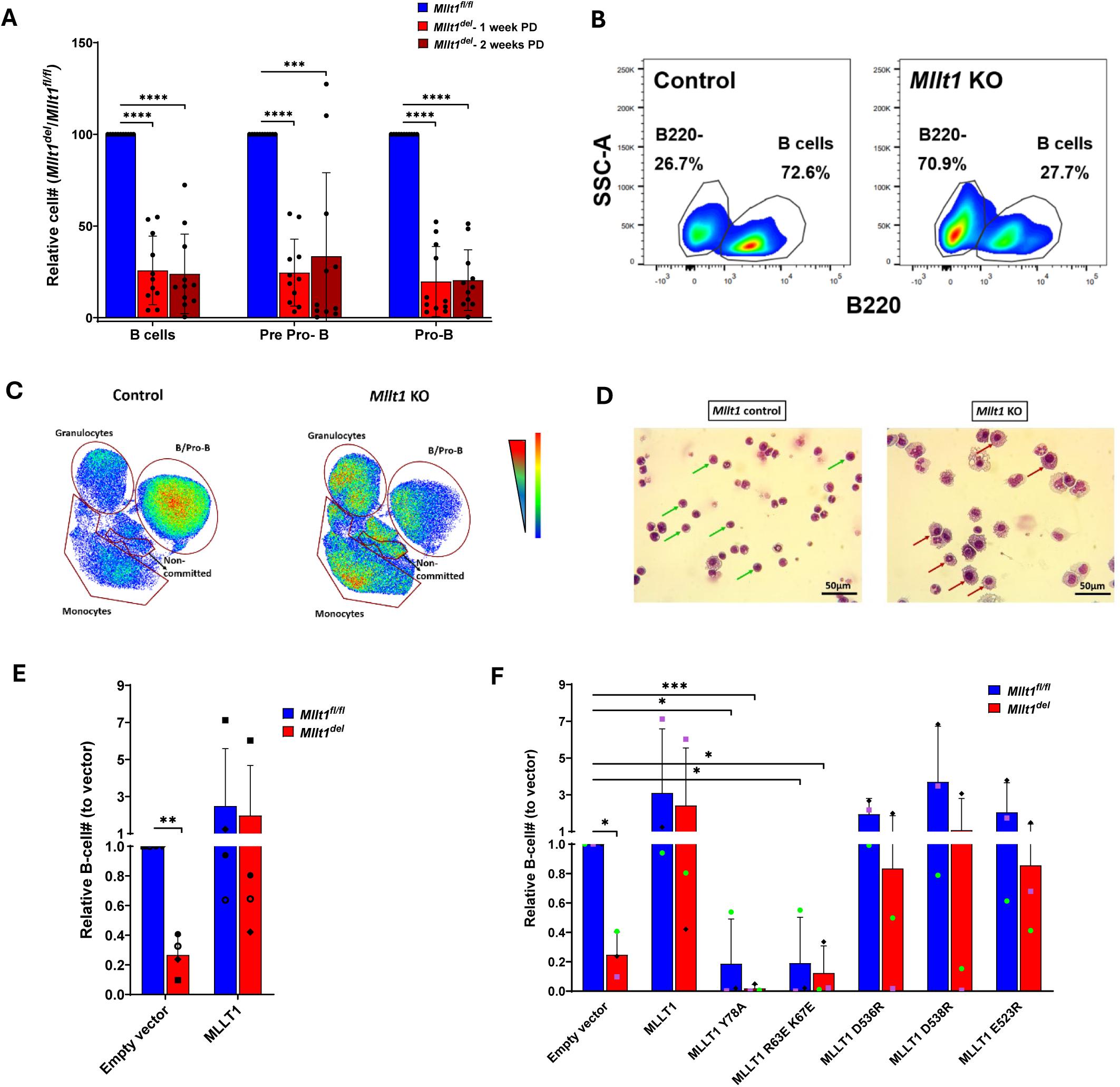
*Mllt1* is Required for B Cell Differentiation *in vitro*. **(A)** Relative number of B cell subpopulations following *in vitro Mllt1* deletion at 1 or 2 weeks post-deletion (PD) (n = 11 mice). **(C)** Representative flow plot of B220 surface expression in vehicle and *Mllt1^del^* B-enriched bone marrow cells following *in vitro* culture for 2 weeks. **(C)** UMAP plot shows different populations of B cells, non-committed cells and myeloid cells following 2 weeks post *in vitro Mllt1* deletion. **(D)** Cytospin Wright-Giemsa staining images indicating lymphoid cells (green arrows) and myeloid cells (red arrows) in *Mllt1^fl/fl^* control and *Mllt1^del^* cells following *in vitro* culture. **(E)** Relative number of B cells in complementation experiments introducing wild type MLLT1 or mCherry vector alone in *in vitro* B cell culture in the presence or absence of endogenous *Mllt1* (n = 4). **(F)** Relative number of B cells following complementation with various MLLT1 mutants lacking specific functions with wild type add-back and vector in the presence or absence of endogenous *Mllt1* (n = 3). Data shown as mean ± SD. *p < 0.05, **p < 0.01, ***p < 0.001, ****p < 0.0001 by One sample t test.

### Complementation with wild type MLLT1, but not YEATS domain mutants, recovers the loss of B cells following Mllt1 deletion

To delineate the specific contribution of Mllt1 in B lineage development, we assessed the ability of exogenous wild type MLLT1 to restore the number of B cells in the absence of endogenous Mllt1. For these experiments, the *in vitro* B cell differentiation experiment was modified to introduce retroviral vectors that express either mCherry alone (vector control) or full length WT MLLT1 in the same vector. As expected, complementation with wild type MLLT1 reestablished the number of B cells and Pro-B cells in contrast to cells complemented with empty vector (Figure 2E; *Online Supplementary Figure S2G*).

Further, to determine which of the known MLLT1 functional domains are necessary for normal B-cell development, we transduced different mutant MLLT1 constructs using the same *in vitro* system. Each of these mutants disrupt a single specific Mllt1 function. In MLLT1 protein, similar to its paralog MLLT3 (AF9), the D536R C-terminal domain point mutant disrupts the binding of MLLT1 to AFF1 (AF4) (a major protein in the transcription super elongation complex SEC); the D538R point mutant interrupts the binding to DOT1L (the H3K79 methyltransferase); and the E523R point mutant prevents the binding of BCOR (a component of non-canonical Polycomb Repressive Complex 1.1) [8, 9, 11, 19, 20]. Similarly, Y78A point mutation disrupts the YEATS domain histone acetyl/crotonyl reader function and the domain double mutant R63E R67E disrupts the RNA binding function while maintaining the domain structure [6, 21, 22].

Complementation with these MLLT1 mutants demonstrated that both YEATS domain mutants failed to restore the loss of B cells and Pro-B cells in the absence of endogenous Mllt1 (Figure 2F; *Online Supplementary Figure S2H*). Further, both YEATS domain mutants had a dominant negative effect with decreased B cells and Pro-B cells, even in the presence of endogenous Mllt1. In contrast, the C-terminal domain mutants largely complemented loss of Mllt1 in the recovery of B and Pro B cells. (Figure 2F; *Online Supplementary Figure S2H*). In conclusion, reintroducing wild type MLLT1 restored B-cell development and the histone acetyl/crotonyl reader function and the RNA binding functions of MLLT1 are essential for this process.

### Mllt1 regulates the expression of B-cell development signature genes

To determine the genes that are regulated by Mllt1 to drive B-cell development, gene expression profiling and ChIP-seq data mining were performed. RNA was isolated from cells following *in vitro* deletion of *Mllt1* and the expression profile was compared with the vehicle control. Gene expression profiling of *Mllt1*-deficient cells by RNA-seq revealed the downregulation of many signature genes of B-cell development (Figure 3A). Gene set enrichment analysis (GSEA) showed downregulation of gene sets associated with B-cell development from both *in vitro*-*Mllt1* deleted cells and B-lineage progenitors of *Mllt1^del^* mice *in vivo* (Figure 3B). Similarly, several B-lineage related gene sets from GSEA of B-lymphoid progenitors were differentially expressed in *Mllt1^del^* mice (Figure 3C). To identify direct targets of Mllt1, we mined previously published ChIP-Seq data of genes directly bound by MLLT1 in a B-cell acute lymphocytic leukemia cell line [23].

**Figure 3.**
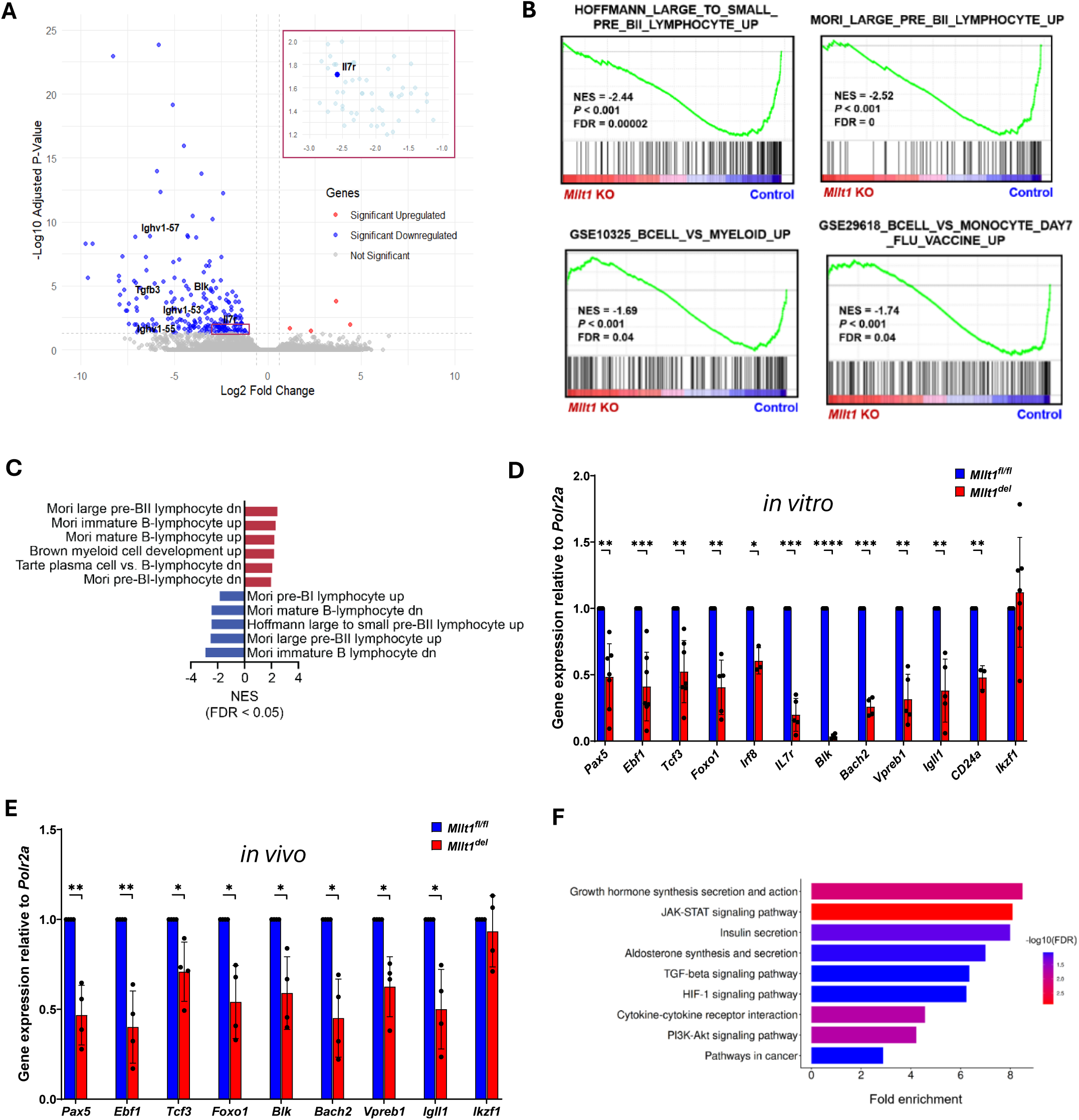
Expression of B Cell Signature Genes is Reduced in the Absence of *Mllt1*. **(A)** RNA sequencing data highlights the differential expression of B-cell development genes upon *Mllt1* deletion *in vitro* when compared to non-deleted vehicle-treated cells (n = 3). **(B)** Gene set enrichment analysis (GSEA) demonstrates the varied pathway gene sets of B-cell development for (top) *Mllt1^del^* cells cultured *in vitro* compared to vehicle (n = 3) and (bottom) B-lymphoid progenitors (B220+ CD43+) isolated from Mllt1^del^ mice compared to Mllt1^fl/fl^ control (n = 3). **(C)** Differentially expressed B-lineage associated gene sets of B-lymphoid progenitors from Mllt1^del^ mice compared to Mllt1^fl/fl^ control by GSEA (n = 3). **(D)** qRT-PCR of cells cultured *in vitro* shows differential expression of B-cell development genes upon loss of *Mllt1* (n ≥ 4). **(E)** qRT-PCR of B-enriched bone marrow cells shows differential expression of early B-cell development genes following the loss of *Mllt1 in vivo* (n = 4). **(F)** Gene ontology analysis of KEGG pathway gene sets for *Mllt1^del^* cells compared to vehicle (n = 3).Data shown as mean ± SD. *p < 0.05, **p < 0.01, ***p < 0.001, ****p < 0.0001 by One sample t test.

Although not highlighted in the publication, we found that key regulatory genes of B cell development, such as *IL7R*, *EBF1*, *PAX5* and *BLK*, were direct MLLT1 targets. In addition, it was previously shown that the master transcription factors Ebf1 and Pax5 act as a part of a B-lineage transcription factor network to directly or indirectly regulate other genes in the network including *Bach2*, *Foxo1*, *Vpreb1* and *Igll1* [24–29]. Using q- RT-PCR, the expression of all these critical B-cell development genes was attenuated by the loss of *Mllt1 in vitro* (Figure 3D). The expression of these same key genes was also significantly decreased in B-enriched lineage negative cells isolated from the bone marrow of *Mllt1*^del^ mice (Figure 3E). Gene ontology analysis revealed the genes involved in the primary signaling pathways of B-cell development such as JAK-STAT and PI3K-Akt signaling are downregulated in the *Mllt1*-deficient *in vitro* cells (Figure 3F). Together, the Mllt1-dependent decrease in the expression of key mediators of early B cell development following *Mllt1* deletion inhibits B cell lineage differentiation.

### Decreased proliferation of developing B cells following Mllt1 deletion

To identity the mechanisms by which *Mllt1* loss reduced the number of B cells, we used Ki67+ staining to investigate the proliferation of B-lineage progenitors *in vivo*.

Proliferation of all the developing B cell populations starting from Early Pre-B were significantly decreased following *Mllt1* deletion (Figure 4A). Gene set enrichment analysis of sorted early B-lineage progenitors revealed negative enrichment of proliferation, cell cycle, DNA replication and Myc related gene sets in the absence of *Mllt1* (Figure 4B). Apart from being the primary receptor essential for B cell fate commitment, IL7 receptor engagement is known to promote proliferation in developing B cells through activation of JAK-STAT and PI3K-AKT signaling pathways [30]. Since *Il7r* gene expression is reduced in *Mllt1*^del^ cells (Figure 3), we determined the surface expression of Il7r in B-enriched lineage negative bone marrow cells. Predictably, the number of surface-Il7r expressing cells were reduced in the B-enriched BM cells isolated from *Mllt1^del^* mice compared to control mice (Figure 4C).

**Figure 4.**
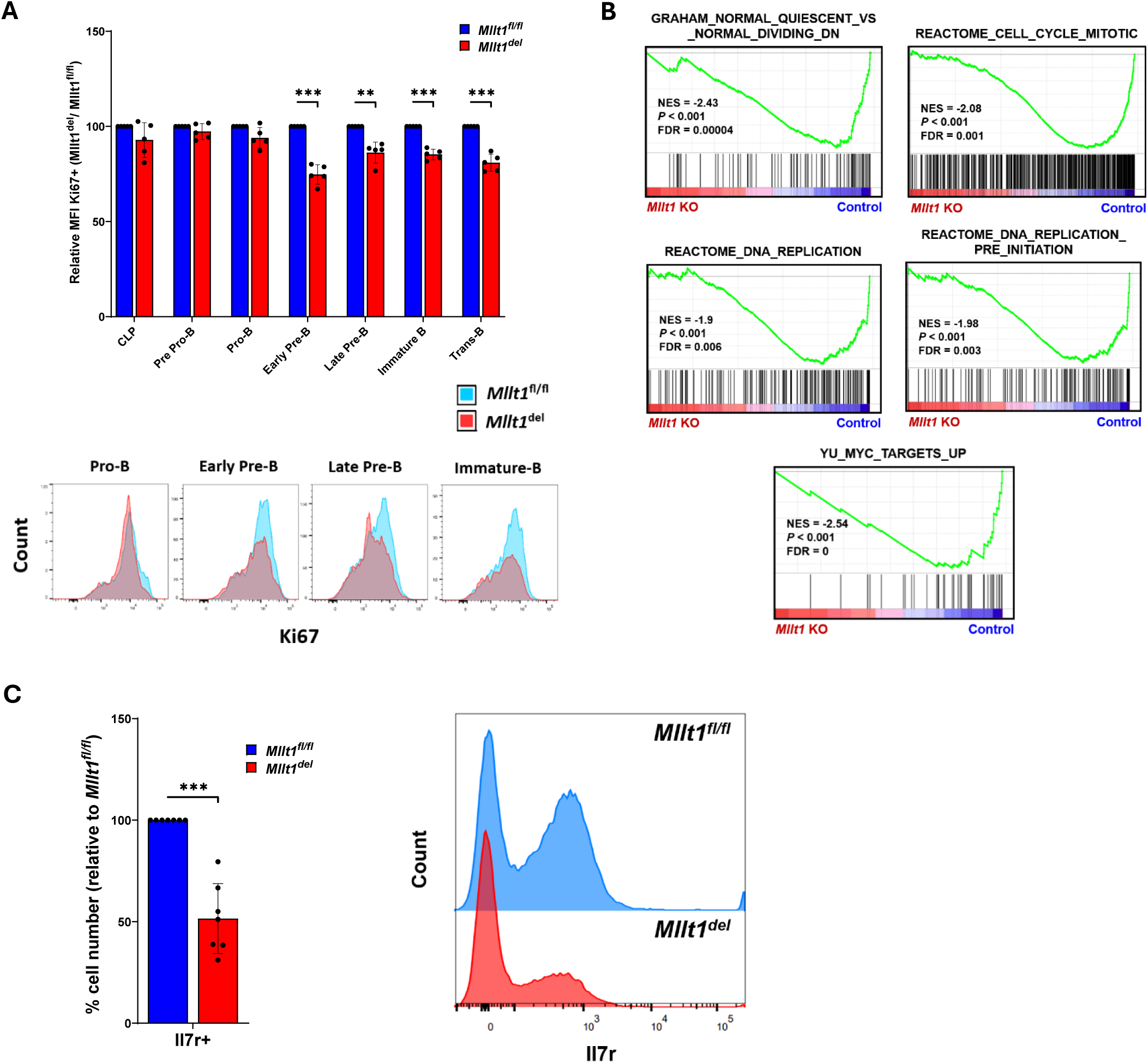
Proliferation of Specific B Cell Subpopulations and Surface Expression of IL7R is Decreased by the Loss of *Mllt1*. **(A)** (*Top*) Cell proliferation measured as mean fluorescence intensity (MFI) of Ki67 staining by intracellular flow cytometry analysis is shown following *in vivo Mllt1* deletion compared to *Mllt1^fl/fl^* control mice (n = 5) and an example of flow plot (*Bottom*). **(B)** Gene set enrichment analysis of RNA-Seq data highlighting cell proliferation-related gene sets for B-lymphoid progenitors (B220+ CD43+) isolated from *Mllt1^del^* mice compared to *Mllt1^fl/fl^* control (n = 3). **(C)** (Left) Relative number of B-lineage enriched bone marrow cells expressing IL7R receptor in *Mllt1^fl/fl^* control and *Mllt1^del^* mice in vivo (n = 7) and an example of flow plot (Right). Data shown as mean ± SD. *p < 0.05, **p < 0.01, ***p < 0.001 by One sample t test.

### Mllt1 deficiency reduces mitochondrial function in progenitor B cells

Mitochondrial activity and membrane potential has been found to increase in differentiating hematopoietic cells to meet energy demands [31, 32]. To assess the effect of *Mllt1* loss on mitochondrial metabolism, measurement of cellular metabolism was performed with an extracellular flux (Seahorse) assay. Lineage positive cells including B-lineage showed decreased OCR (Oxygen Consumption Rate), maximal respiration and spare respiratory capacity following *Mllt1* deletion (Figure 5A). Gene set enrichment analysis of RNAseq data demonstrated that mitochondrial pathways are negatively enriched for *Mllt1*^del^ cells compared to control (Figure 5B). Following the reduction of mitochondrial pathway gene sets, we investigated the effect of loss of *Mllt1* on mitochondrial function of developing B cells. *S*taining of mitochondrial mass and function using MTG and MTR mitotracker dyes showed that Pro-B cells of *Mllt1^del^* mice displayed decreased mitochondrial function over mass compared to controls; however, neither mass nor function were affected in Pre Pro-B cells (Figures 5C-D). This decrease in mitochondrial function in Pro-B but not Pre Pro-B of *Mllt1^del^* mice correlates with the profound defect we observe at the Pre Pro-B to Pro-B transition. These findings indicate that mitochondrial function plays a key role at the differentiation point of Pre Pro-B to Pro-B stage.

**Figure 5.**
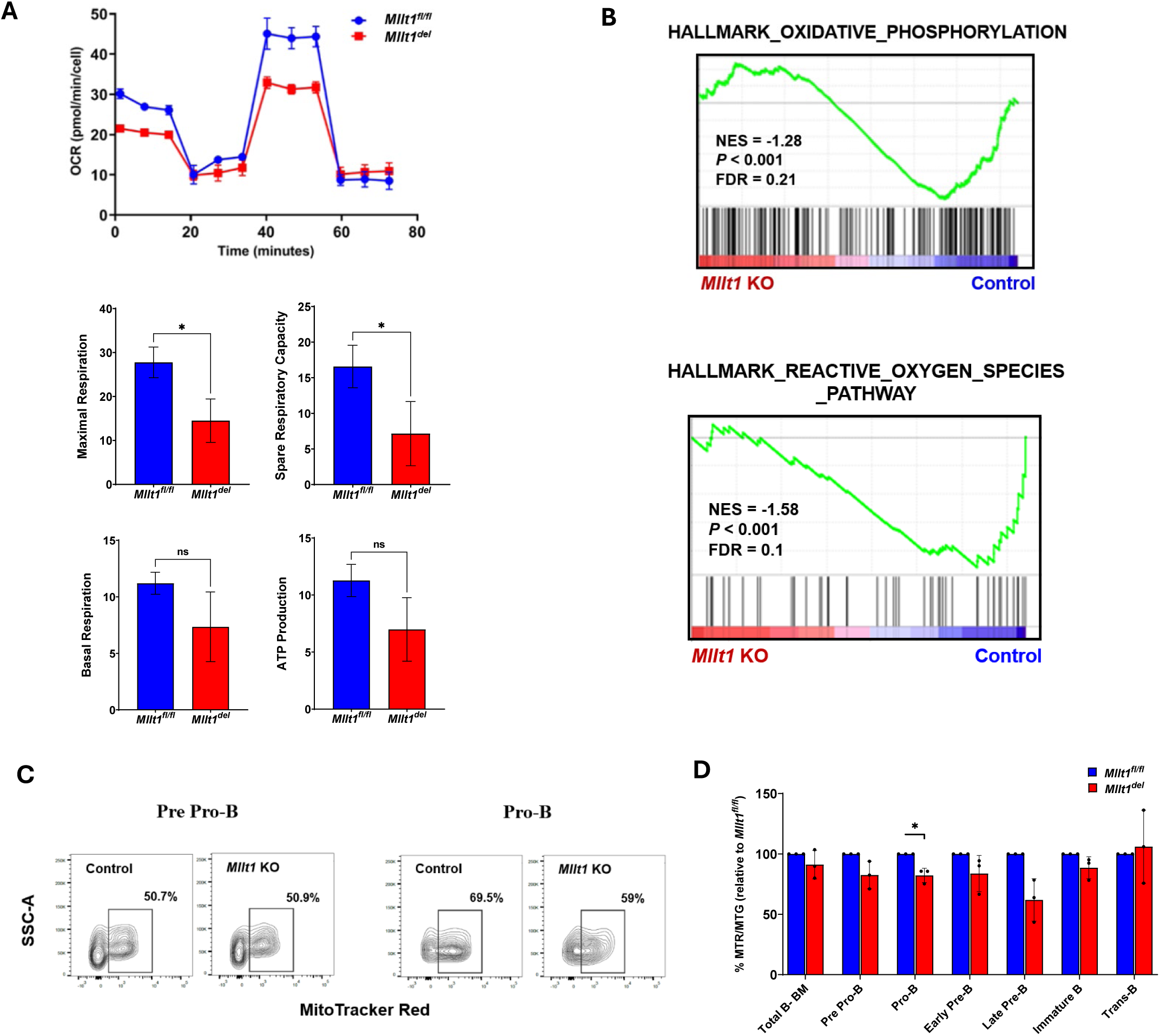
Mitochondrial Function is Reduced in Specific Bone Marrow B Cell Subpopulations Following *Mllt1* Deletion. **(A)** (Top) Seahorse Mito Stress Test of Lineage positive bone marrow cells show oxygen consumption rate plot of *Mllt1^del^* mouse compared to *Mllt1^fl/fl^* control. (Bottom) Quantification of seahorse assay parameters including maximal respiration and spare respiratory capacity. **(B)** Gene set enrichment analysis shows the differentially enriched mitochondrial pathway gene sets between *Mllt1^del^* and non-deleted *Mllt1 in vitro* cultured cells (n = 3). **(C)** Flow cytometry plot using MitoTracker Red (MTR) to identify the function (membrane potential) of mitochondria highlights the difference in mitochondrial function between *Mllt1^fl/fl^* control and *Mllt1^del^* mice. **(D)** Relative ratio of MTR over MTG (MitoTracker Green to identify mitochondrial mass) for developing B cells of bone marrow following *Mllt1* deletion compared to control mice (n = 3). Data shown as mean ± SD. *p < 0.05 by One sample t test.

## DISCUSSION

Our study demonstrates the critical cell-intrinsic role of Mllt1, a chromatin reader protein, in the B lymphocyte lineage. Conditional *Mllt1* deletion significantly blocked B lineage development through disrupted expression of critical lineage specification transcription factors and signaling pathways. Furthermore, effects on mitochondrial function, cell proliferation and B cell survival signals contribute to the observed phenotype. This study determines a novel functional role and mechanism of Mllt1 in regulating B lymphopoiesis (*Online Supplementary Figure S4*).

### Conditional deletion of *Mllt1* arrests B cell development

The conditional deletion of *Mllt1* in our model *Mllt1^fl/fl^; Rosa26CreER^T2/+^* mice arrested B cell development at the Pre Pro-B stage, with developing B cells from the Pro-B stage to transitional B cells dramatically reduced. It is striking that no other bone marrow cell populations were significantly affected by *Mllt1* loss. Analysis of secondary lymphoid organs revealed a specific decrease of spleen transitional B cells, immature B cells which constitute about 10% of the splenic B cell population [33]. In contrast to mature splenic B cells which were not affected, recirculating B cells in lymph nodes and peripheral blood were decreased following *Mllt1* deletion. Significantly reduced *Baffr* expression, which encodes the receptor for B cell survival in peripheral blood [15, 16], may explain the observed B-lymphopenia, while decreased bone marrow *S1pr1* expression, important for egress [17, 18], may contribute to reduced spleen transitional B cells. The B-lineage phenotype following deletion of *Mllt1* was recovered at later time points, likely due to restored B-cell homeostasis under steady-state conditions to maintain a constant number of B cells [34, 35]. Investigation of B cell differentiation *in vitro* confirmed the cell intrinsic requirement of *Mllt1* for B cell differentiation without an effect on myeloid cells. Interestingly, the observed B cell phenotype of *Mllt1* deletion contrasts with its very close paralog, Mllt3. Although Mllt3 has very similar functional domains as Mllt1, it is important for maintenance of LT-HSC function [12]. This may be due to differential expression pattern of Mllt3 in normal hematopoietic cells compared to Mllt1.

### Functional MLLT1 YEATS domain required for B cell differentiation

MLLT1 complementation in the absence of endogenous mouse Mllt1 was able to recover the B cell phenotype. However, complementation with MLLT1 YEATS domain mutants lacking either the acetyl/crotonyl histone tail reader or the RNA binding functions [4–6, 21, 22] did not restore the B-cell phenotype and further had a dominant negative effect in the presence of endogenous Mllt1. Mllt3 condensate formation is dependent on YEATS domain-RNA binding [6], so it is plausible that RNA dependent condensate formation of Mllt1 contributes to B cell differentiation. However, further studies will be needed to elucidate the mechanisms underlying these observations.

### Mllt1 regulates B-lineage specification factors and signaling pathways

Mllt1 deletion resulted in decreased expression of B cell related genes, as well as developmental and critical signaling pathways of early B-cell development [36]. The transitional block at Pre Pro-B stage in *Mllt1*-deficient mice replicates the phenotype observed following the loss of critical B-lineage specification transcription factors in mice, Ebf1 [37] and E2A [38]. Several key genes of B-lineage commitment, including *EBF1*, *PAX5* and *BLK* are direct targets of MLLT1 [23] and had decreased expression following both *in vivo* and *in vitro Mllt1* deletion. Expression of additional key genes of B-lineage commitment and differentiation, including *E2A (Tcf3), Foxo1, Bach2*, *Vpreb1* and *Igll1*, were decreased following *Mllt1* depletion. Some of these genes, *Bach2*, *Vpreb1* and *Igll1*, are direct targets of Pax5 signaling [39–41]. B-lineage specification, commitment and differentiation is the result of a combinatorial effect of transcription factor networks through hierarchical forward steps and feedback loops. Therefore, the decrease of developing B cells is likely due to the combination of inter-regulatory effects of the above-mentioned transcription factors, in addition to the effects on direct target genes, following Mllt1 loss. Apart from these transcription factors, the surface expression of a critical signaling receptor for B-cell development, Il7r, was reduced following both *in vivo* and *in vitro* deletion of *Mllt1*. These results support the requirement of Mllt1 for maintenance of the B-lineage transcription network and Il7r signaling pathway.

### Mllt1-dependent effects on proliferation and mitochondrial function

Further, exploring the functional consequences at the cellular level revealed that proliferation and mitochondrial function of developing B cells in the bone marrow were reduced with the loss of *Mllt1*. Cell proliferation was significantly decreased in all stages of B-lineage progenitors post Pro-B stage, correlating with the observed reduction in the number of respective cells. Notably, the extent of this decrease was higher in Early Pre-B, the most proliferative stage of developing B cells as they undergo clonal expansion after the successful rearrangement of the immunoglobulin heavy chain (IgH) and subsequent expression of the pre-BCR [42, 43]. Since Il7r signaling and the master B-lineage transcription factors Ebf1, Pax5 and Foxo1 are critical regulators of IgH rearrangement [37, 44–46], the reduced proliferation of Early Pre-B could be the direct outcome of the reduced expression of Il7r and the B-lineage transcription factors following the loss of Mllt1. Together, these results suggest that the dramatic decrease in proliferation of Pre-B cells is the result of a deficiency in the pre-BCR mediated proliferation signal. In stem cells, it has been shown that increased mitochondrial activity aids cell proliferation by providing sustained ATP as energy source and metabolites as signaling molecules [47]. Thus, extrapolating this connection to other bone marrow progenitor cells, the reduced mitochondrial function as a result of Mllt1 loss contributes to the reduced proliferation and resulting decrease of B cells.

Our results identify the novel role of Mllt1 in regulating early B-cell development. In addition to its normal role in B cell lymphopoiesis, our data suggest a plausible role in KMT2A-MLLT1 leukemia, which is predominantly mixed phenotype or B lineage-ALL. As a result of the chromosomal translocation to create the oncogene driver in this leukemia subtype, one wildtype MLLT1 allele is lost, blocking efficient B lineage differentiation.

Further experiments to determine how this may contribute toward the resulting B-lineage phenotype are warranted.

## Acknowledgments

This work was supported by NIH R01CA233749 awarded to NJZ, JHB, MEF and CSH. We would like to acknowledge the Loyola Flow Cytometry Core Facility (RRID:SCR_025109), especially Robert Ladd, for his assistance with flow cytometry and sorting.

## Data availability

The RNAseq data have been deposited at GEO under superseries GSE342909.

## Notes

https://www.ncbi.nlm.nih.gov/geo/query/acc.cgi?acc=GSE342909

### Competing Interest Statement

The authors have declared no competing interest.

