## Supplemental Figures and Methods for "The chromatin reader protein MLLT1 is critical to maintain normal B lymphopoiesis"

#### Supplementary Figure Legends

**Figure S1. Number of B Cells is Recovered *in vivo* in Most Tissues at 5 Weeks Post *Mllt1* Deletion. (A)** Plot based on surface marker flow cytometry analysis represents the total number of B cells, B progenitors and mature B cells from the hind limbs of $Mllt1^{fl/fl}$ control and $Mllt1^{del}$ mice (n = 7). **(B)** Spleen weight to body weight ratio of $Mllt1^{fl/fl}$ control and $Mllt1^{del}$ mice (n = 7). **(C)** Concentration of different immunoglobulin levels in mice plasma at 2 weeks post *Mllt1* deletion (n = 3). **(D)** Relative number of different bone marrow cell populations following *in vivo Mllt1* deletion (n ≥ 2). **(E)** Relative number of developing B cells of bone marrow (n = 6), spleen B-cell subpopulations (n = 3), B cells of inguinal lymph nodes (n = 3) and recirculating B cells of peripheral blood (n = 8) following 5 weeks post *in vivo Mllt1* deletion. **(F)** Relative ratio of MTR (MitoTracker Red) over MTG (MitoTracker Green) for developing B cells of bone marrow at 5 weeks following *Mllt1* deletion compared to control mice (n = 3). **(G)** MFI of Ki67 staining between $Mllt1^{fl/fl}$ control and $Mllt1^{del}$ mice 5 weeks post deletion. Data shown as mean ± SD. *p < 0.05, ***p < 0.001 by an unpaired Student t test for (A) and (C) and by One sample t test for (B), (D), (E), (F) and (G).

**Figure S2. *Mllt1* Regulates B Cell Differentiation *in vitro*. (A)** Total number of cells and B lineage subpopulations following *in vitro Mllt1* deletion (n = 11). **(B)** Representative flow plots of surface expression of myeloid cell markers CD115, CD11b and Gr-1 in $Mllt1^{del}$ samples compared to vehicle. **(C)** Total cell numbers of B cell and myeloid cells following *in vitro Mllt1*-deletion. Colors indicate paired samples (n = 7-10 mice). **(D)** Relative number of myeloid lineage cell subpopulations following *in vitro Mllt1* deletion (n = 7-10 mice). **(E)** Relative number of B and Pro-B cells from $Mllt1^{fl/fl}$ cells treated *in vitro* with 4-hydroxy tamoxifen compared to treatment with vehicle ethanol (n = 3). **(F)** Relative number of B and Pro-B cells following *in vitro* deletion of one *Mllt1* allele (n ≥ 2). **(G)** Relative number of Pro-B cells in complementation experiments introducing wild type MLLT1 or mCherry vector in *in vitro*

**Figure S3. Loss of *Mllt1* Decreases the Expression of *Baffr* in the Bone Marrow and Peripheral Blood and Expression of *S1pr1r* in the Bone Marrow. (A)** qRT-PCR of *Baffr* from B-enriched lineage negative bone marrow and spleen cells, and non-enriched peripheral blood cells following *Mllt1* deletion (n = 4-6). **(B)** qRT-PCR of *S1pr1* from B-enriched lineage negative bone marrow and spleen cells following *Mllt1* deletion (n = 4). Data shown as mean ± SD. *p < 0.05 by One sample t test.

**Figure S4. Mllt1 is Required for B Cell Development and Survival.** Loss of Mllt1 reduces the number of developing B cells in bone marrow, transitional B cells in spleen, and mature recirculating B cells in lymph nodes and peripheral blood. Mechanistically, Mllt1-deficiency induced B cell reduction is due to decrease in the expression of B-lineage target genes, mitochondrial function, proliferation of developing B cells and finally by downregulation of the expression of *Baffr*, a critical gene for B-cell survival.

Complementation with wild type *MLLT1* restores the number of B cells *in vitro* in the absence of endogenous Mllt1 but not the YEATS domain mutants lacking reader function and RNA binding function. Created in BioRender. Prakash, J. (2026) https://BioRender.com/6se73n1.

### Online Supplementary Figures:

#### Figure S1

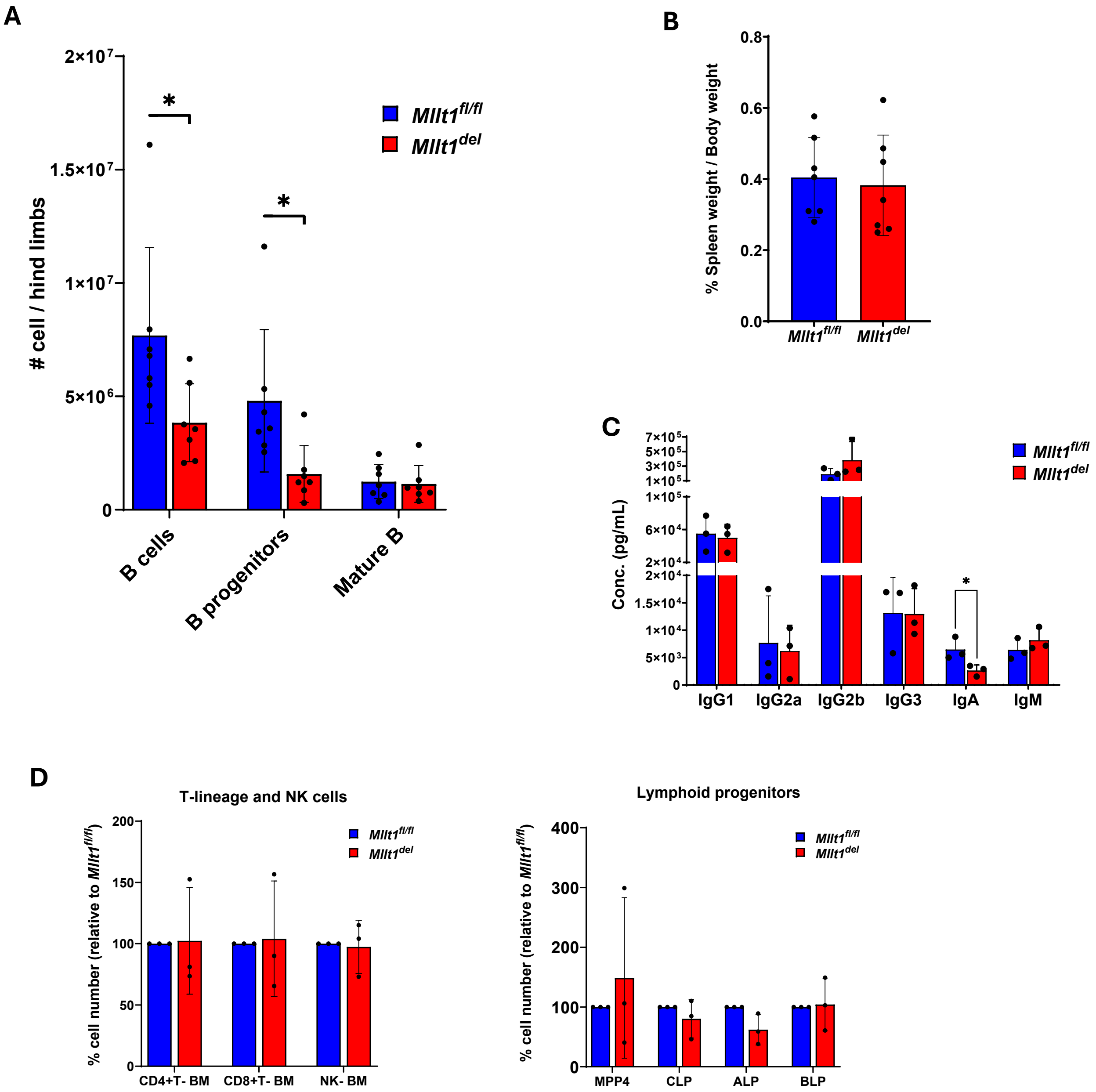

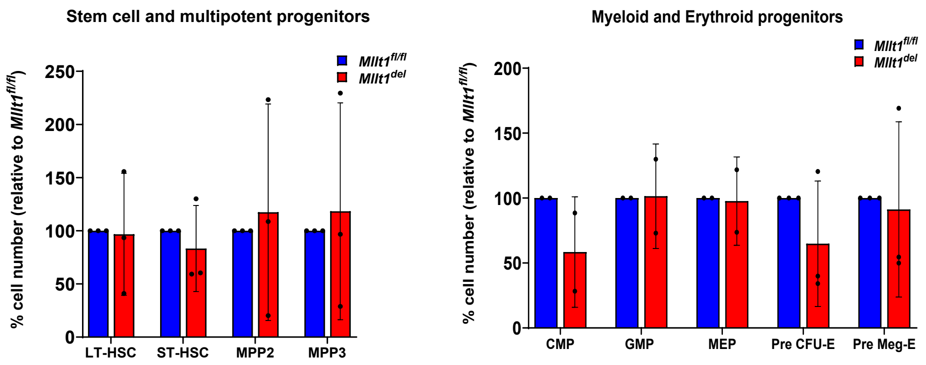
Stem cell and multipotent progenitors
% cell number (relative to Mllt1fl/fl)
Mllt1fl/fl
Mllt1del
LT-HSC
ST-HSC
MPP2
MPP3
Myeloid and Erythroid progenitors
CMP
GMP
MEP
Pre CFU-E
Pre Meg-E

E

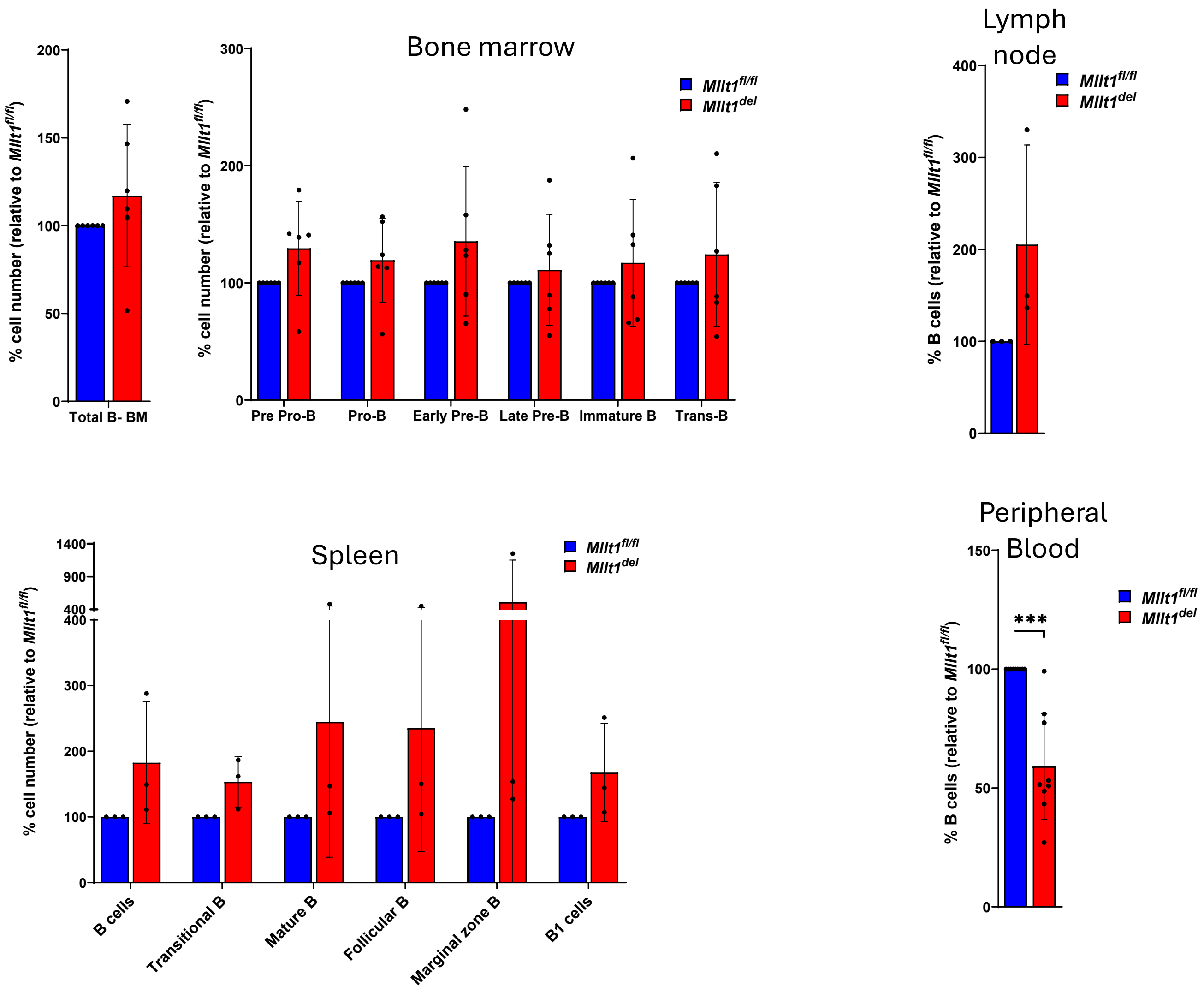
% cell number (relative to Mllt1fl/fl)
Total B- BM
Bone marrow
Mllt1fl/fl
Mllt1del
Pre Pro-B
Pro-B
Early Pre-B
Late Pre-B
Immature B
Trans-B
Lymph node
% B cells (relative to Mllt1fl/fl)
Spleen
B cells
Transitional B
Mature B
Follicular B
Marginal zone B
B1 cells
Peripheral Blood
***

F

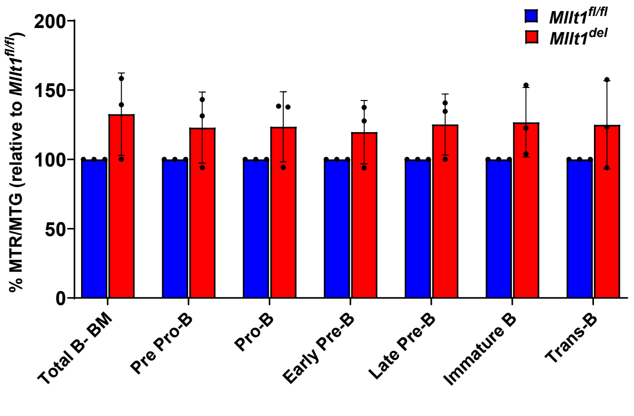
Mllt1fl/fl
Mllt1del
% MTR/MTG (relative to Mllt1fl/fl)
200
150
100
50
0
Total B- BM
Pre Pro-B
Pro-B
Early Pre-B
Late Pre-B
Immature B
Trans-B

G

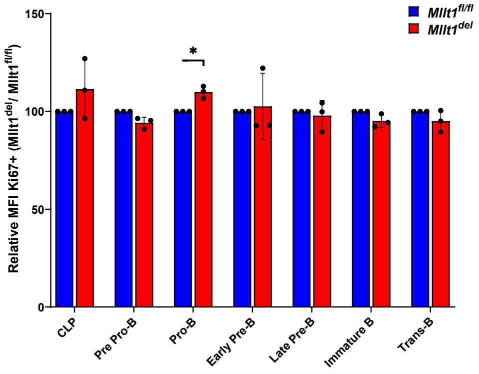
Mllt1fl/fl
Mllt1del
Relative MFI Ki67+ (Mllt1del/ Mllt1fl/fl)
150
100
50
0
*
CLP
Pre Pro-B
Pro-B
Early Pre-B
Late Pre-B
Immature B
Trans-B

### Figure S2

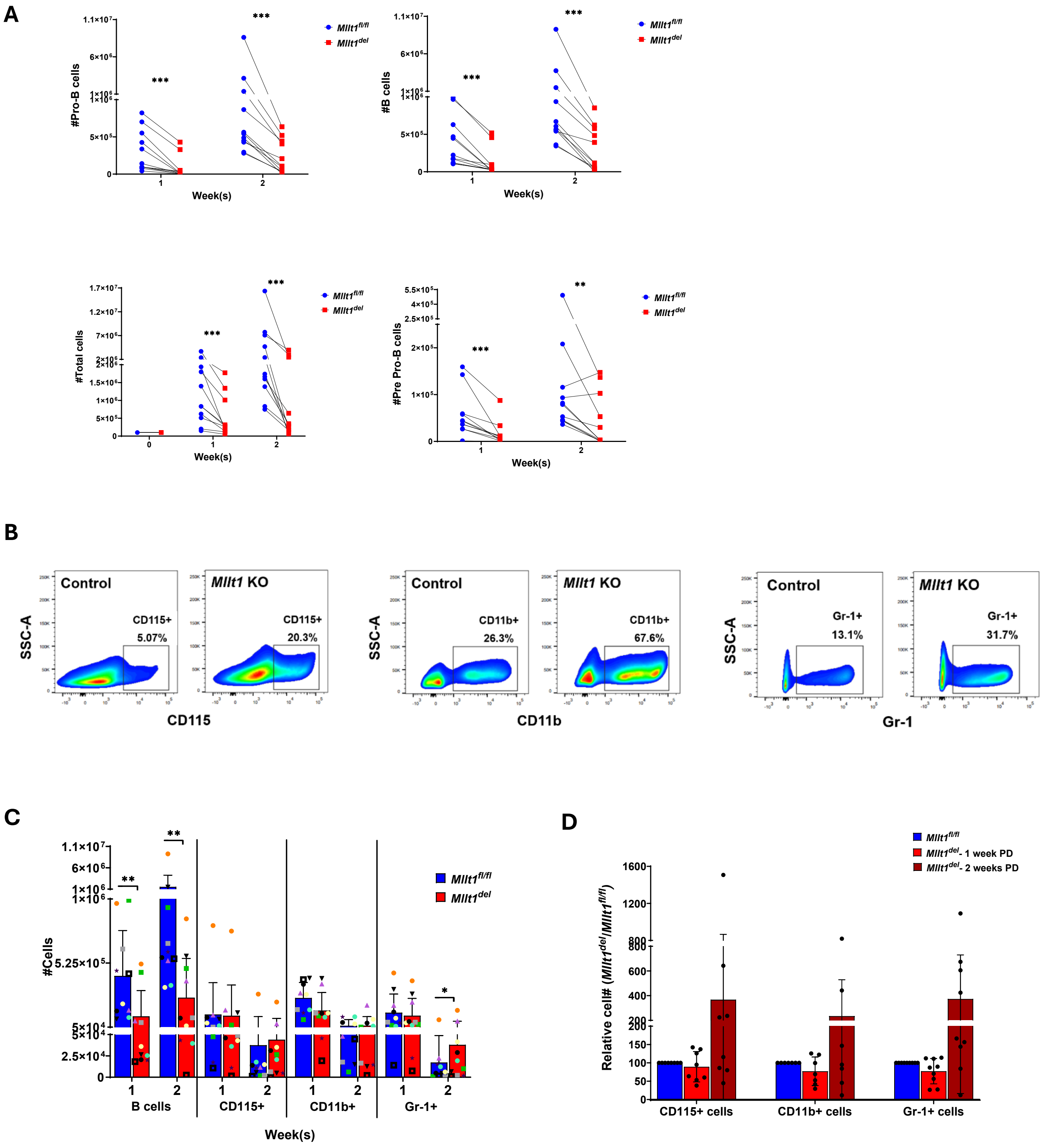
A
#Pro-B cells
Week(s)
#B cells
#Total cells
#Pre Pro-B cells
Mllt1fl/fl
Mllt1del
B
Control
Mllt1 KO
SSC-A
CD115
CD115+
5.07%
20.3%
CD11b
CD11b+
26.3%
67.6%
Gr-1
Gr-1+
13.1%
31.7%
C
#Cells
B cells
CD115+
CD11b+
Gr-1+
Week(s)
D
Relative cell# (Mllt1del/Mllt1fl/fl)
Mllt1del- 1 week PD
Mllt1del- 2 weeks PD
CD115+ cells
CD11b+ cells
Gr-1+ cells

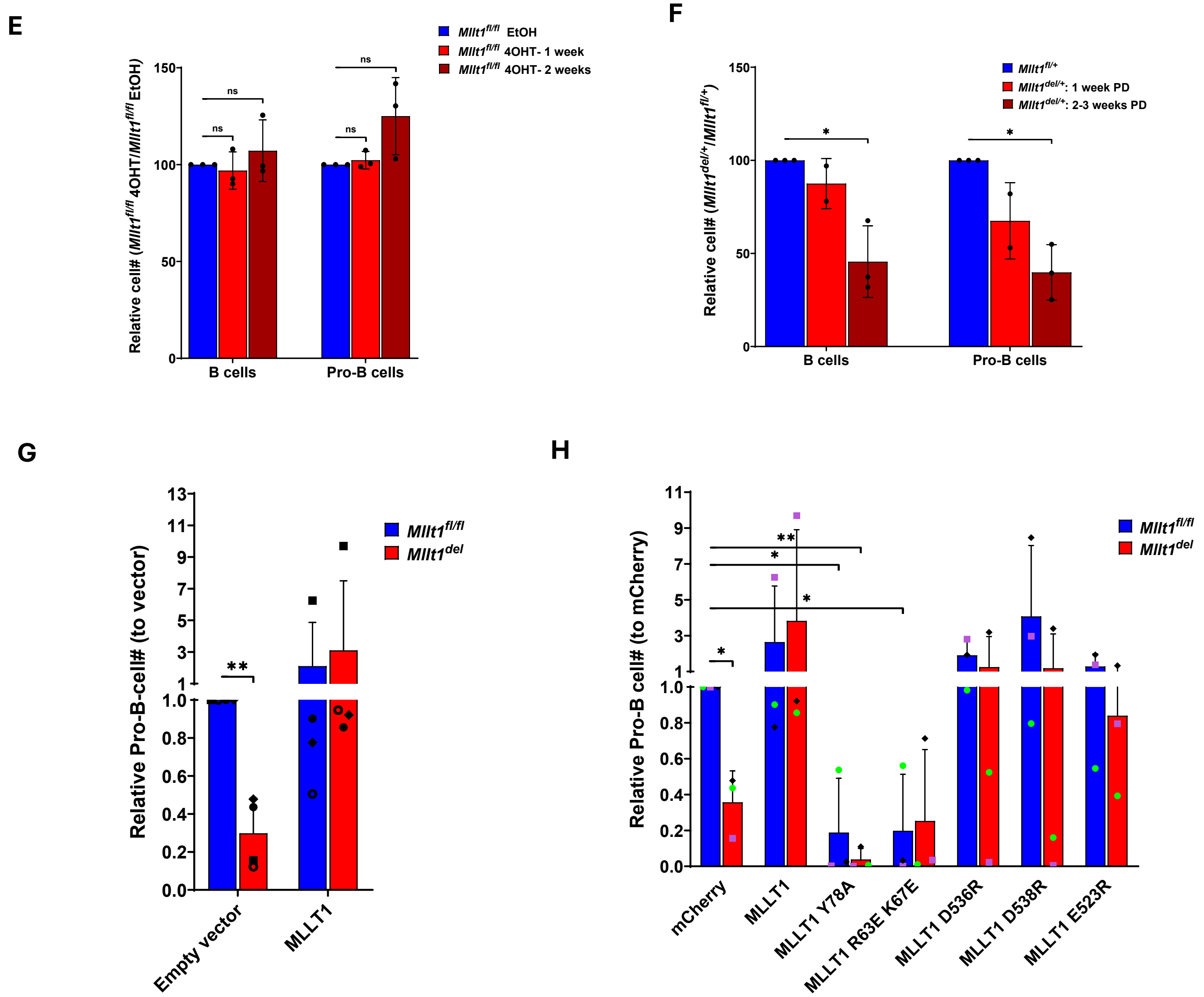

E
Mllt1fl/fl EtOH
Mllt1fl/fl 4OHT- 1 week
Mllt1fl/fl 4OHT- 2 weeks
Relative cell# (Mllt1fl/fl 4OHT/Mllt1fl/fl EtOH)
150
100
50
0
ns
B cells
Pro-B cells
F
Mllt1fl/+
Mllt1del/+: 1 week PD
Mllt1del/+: 2-3 weeks PD
Relative cell# (Mllt1del/+/Mllt1fl/+)
150
100
50
0
*
B cells
Pro-B cells
G
Mllt1fl/fl
Mllt1del
Relative Pro-B-cell# (to vector)
13
11
9
7
5
3
1
1.0
0.8
0.6
0.4
0.2
0.0
**
Empty vector
MLLT1
H
Mllt1fl/fl
Mllt1del
Relative Pro-B cell# (to mCherry)
11
9
7
5
3
1
1.0
0.8
0.6
0.4
0.2
0.0
*
**
mCherry
MLLT1
MLLT1 Y78A
MLLT1 R63E K67E
MLLT1 D536R
MLLT1 D538R
MLLT1 E523R

### Figure S3

**A**

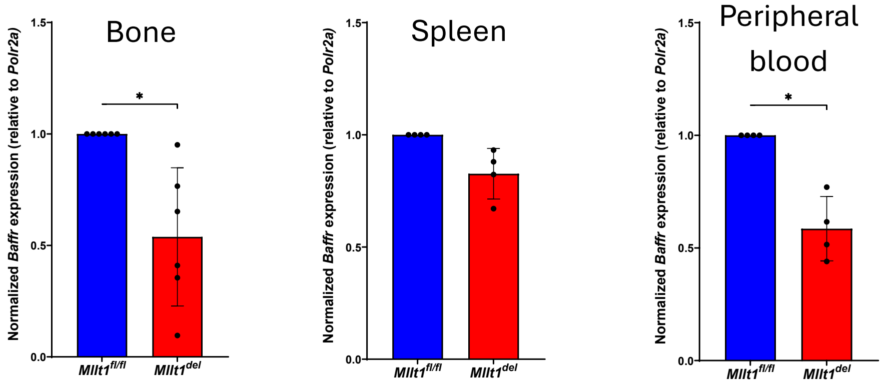
Bone
Spleen
Peripheral blood
Normalized Baffr expression (relative to Polr2a)
1.5
1.0
0.5
0.0
*
Mllt1fl/fl
Mllt1del

**B**

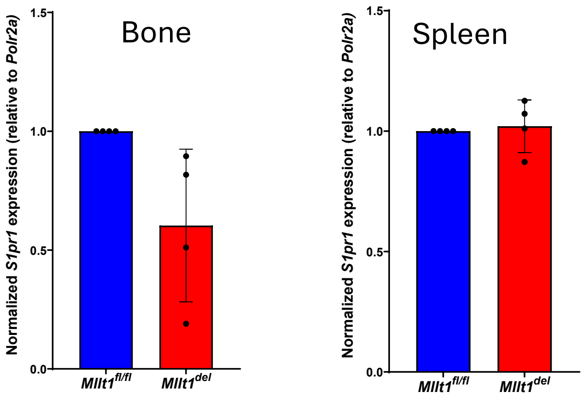
Bone
Spleen
Normalized S1pr1 expression (relative to Polr2a)
1.5
1.0
0.5
0.0
Mllt1fl/fl
Mllt1del

**Figure S4**

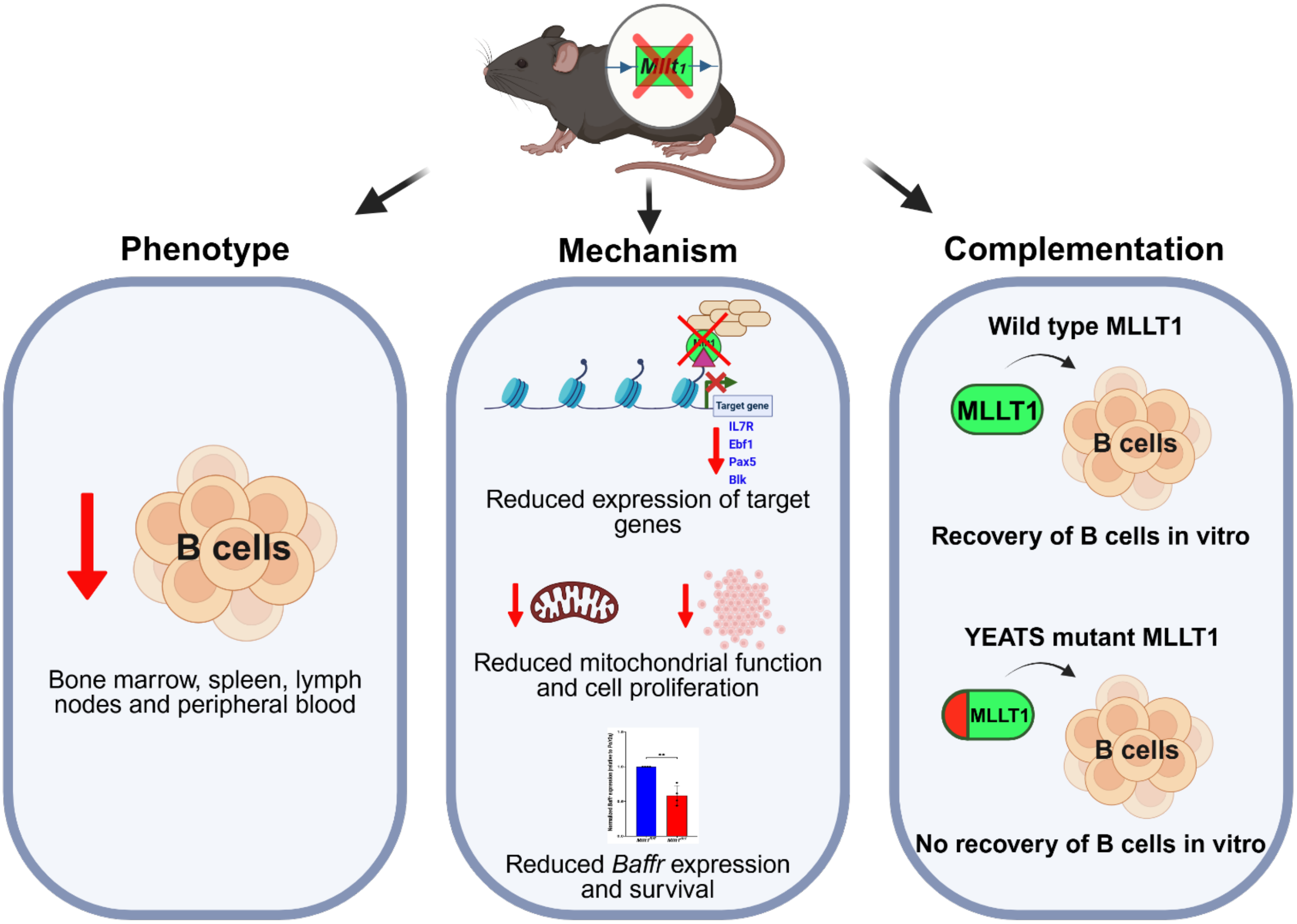

Mllt1
Phenotype
Mechanism
Complementation
B cells
Bone marrow, spleen, lymph nodes and peripheral blood
Target gene
IL7R
Ebf1
Pax5
Blk
Reduced expression of target genes
Reduced mitochondrial function and cell proliferation
Reduced Baffr expression and survival
Wild type MLLT1
MLLT1
B cells
Recovery of B cells in vitro
YEATS mutant MLLT1
MLLT1
B cells
No recovery of B cells in vitro

***Table S1.*** Mllt1 genotyping primers

| **Primer Name** | **Nucleotide sequence (5'– 3')** |
|---|---|
| 1. CSD-Mllt1-F | ATTCGGAGATGTCTTACTGGTCTTGC |
| 2. CSD-Mllt1-ttR | CCAAGTTACTCAGACACAACTGCCTG |
| 3. Cre wt Fwd | AAAGTCGCTCTGAGTTGTTAT |
| 4. Cre wt Rev | GGAGCGGGAGAAATGGATATG |
| 5. Cre mut Rev | CCTGATCCTGGCAATTTCG |

| **Primer Combinations** | **Band size (bp)** | **Genotype** |
|---|---|---|
| 1 & 2 | 324 | Wildtype *Mllt1 (+/+)* |
| 1 & 2 | 448 | Floxed *Mllt1 (fl/fl)* |
| 1 & 2 | 324 + 448 | *fl*/+ |
| 3, 4 & 5 | 825 | *cre/cre* |
| 3, 4 & 5 | 650 + 825 | *cre*/+ |
| 3, 4 & 5 | 650 | +/+ |

***Table S2.*** List of Fluorescent Labeled Antibodies and cell surface markers used for immunophenotyping *in vivo*

**Bone marrow**

**Stem cell and multipotent progenitors**

| Antibody | Fluorophore | Clone | Catalog# & source |
|---|---|---|---|
| Sca-1 | BV510 | D7 | 108129 – Biolegend |
| c-Kit | APC | 2B8 | 105812 – Biolegend |
| Flk2 | PerCP-eFluor 710 | A2F10 | 46-1351-82 – eBioscience (Invitrogen) |
| CD48 | APC-Cy7 | HM48-1 | 103432 – Biolegend |
| CD150 | PE-Cy7 | mShad150 | 25-1502-80 – eBioscience (Invitrogen) |
| Lineage | | | |
| CD3 | BV421 | 145-2C11 | 100335 – Biolegend |
| CD8a | BV421 | 53-6.7 | 100738 – Biolegend |
| Gr1 | BV421 | RB6-8C5 | 108433 – Biolegend |
| Ter119 | BV421 | TER-119 | 116234 – Biolegend |
| B220 | BV421 | RA3-6B2 | 103240 – Biolegend |

**Myeloid progenitors**

| Antibody | Fluorophore | Clone | Catalog# & source |
|---|---|---|---|
| Sca-1 | BV510 | D7 | 108129 – Biolegend |

| | | | |
|---|---|---|---|
| c-Kit | APC | 2B8 | 105812 – Biolegend |
| CD34 | Biotin | RAM34 | 13-0341-85 – eBioscience (Invitrogen) |
| CD16/CD32 | PE-Cy7 | 93 | 101318 – Biolegend |
| Lineage | | | |
| CD3 | BV421 | 145-2C11 | 100335 – Biolegend |
| CD8a | BV421 | 53-6.7 | 100738 – Biolegend |
| Gr1 | BV421 | RB6-8C5 | 108433 – Biolegend |
| Ter119 | BV421 | TER-119 | 116234 – Biolegend |
| B220 | BV421 | RA3-6B2 | 103240 – Biolegend |

**Erythroid progenitors**

| Antibody | Fluorophore | Clone | Catalog# & source |
|---|---|---|---|
| Sca-1 | BV510 | D7 | 108129 – Biolegend |
| c-Kit | APC | 2B8 | 105812 – Biolegend |
| CD41 | PerCP-Cy5.5 | MWReg30 | 133918 – Biolegend |
| CD150 | Biotin | 9D1 | 13-1501-82 – eBioscience (Invitrogen) |
| CD16/CD32 | PE-Cy7 | 93 | 101318 – Biolegend |
| CD105 | BV786 | MJ7/18 | 564746 – BD Horizon |
| Lineage | | | |
| CD3 | BV421 | 145-2C11 | 100335 – Biolegend |

| | | | |
|---|---|---|---|
| CD8a | BV421 | 53-6.7 | 100738 – Biolegend |
| Gr1 | BV421 | RB6-8C5 | 108433 – Biolegend |
| Ter119 | BV421 | TER-119 | 116234 – Biolegend |
| B220 | BV421 | RA3-6B2 | 103240 – Biolegend |

**Lymphoid progenitors**

| Antibody | Fluorophore | Clone | Catalog# & source |
|---|---|---|---|
| Sca-1 | BV510 | D7 | 108129 – Biolegend |
| c-Kit | APC | 2B8 | 105812 – Biolegend |
| IL7R | PE-Cy7 | A7R34 | 135014 – Biolegend |
| Ly6D | Biotin | REA906 | 130-115-310 – Miltenyi Biotec |
| Lineage | | | |
| CD3 | BV421 | 145-2C11 | 100335 – Biolegend |
| CD8a | BV421 | 53-6.7 | 100738 – Biolegend |
| Gr1 | BV421 | RB6-8C5 | 108433 – Biolegend |
| Ter119 | BV421 | TER-119 | 116234 – Biolegend |
| B220 | BV421 | RA3-6B2 | 103240 – Biolegend |

**T and NK cells**

| Antibody | Fluorophore | Clone | Catalog# & source |
|---|---|---|---|
| CD3 | PE-Cy7 | 145-2C11 | 100320 – Biolegend |

| | | | |
|---|---|---|---|
| CD4 | APC | GK1.5 | 100412 – Biolegend |
| CD8 | APC-Cy7 | 53-6.7 | 100714 – Biolegend |
| NK1.1 | PerCP-Cy5.5 | PK136 | 108728 – Biolegend |

**B cell**

| **Antibody** | **Fluorophore** | **Clone** | **Catalog# & source** |
|---|---|---|---|
| B220 | PerCP-Cy5.5 | RA3-6B2 | 103236 – Biolegend |
| CD19 | BV510 | 6D5 | 115546 – Biolegend |
| IgM | BV786 | II/41 | 743328 - BD OptiBuild |
| IgD | BV650 | 11-26c.2a | 405721 – Biolegend |
| CD43 | APC-Cy7 | 1B11 | 121220 – Biolegend |
| CD93 | PE-Cy7 | AA4.1 | 136506 – Biolegend |
| CD24 | APC | 30-F1 | 138506 – Biolegend |
| Dump channel | | | |
| CD3 | BV421 | 145-2C11 | 100335 – Biolegend |
| CD11b | BV421 | M1/70 | 101251 – Biolegend |
| Gr1 | BV421 | RB6-8C5 | 108433 – Biolegend |
| Ter119 | BV421 | TER-119 | 116234 – Biolegend |

**Spleen**

| Antibody | Fluorophore | Clone | Catalog# & source |
|---|---|---|---|
| B220 | PerCP-Cy5.5 | RA3-6B2 | 103236 – Biolegend |
| CD19 | BV510 | 6D5 | 115546 – Biolegend |
| IgM | BV786 | II/41 | 743328 - BD OptiBuild |
| IgD | BV650 | 11-26c.2a | 405721 – Biolegend |
| CD43 | APC-Cy7 | 1B11 | 121220 – Biolegend |
| CD93 | PE-Cy7 | AA4.1 | 136506 – Biolegend |
| CD23 | PE | B3B4 | 101608 – Biolegend |
| CD21/35 | APC | 7E9 | 123412 – Biolegend |

**Peripheral blood & Lymph nodes**

| Antibody | Fluorophore | Clone | Catalog# & source |
|---|---|---|---|
| B220 | PerCP-Cy5.5 | RA3-6B2 | 103236 – Biolegend |
| CD19 | BV510 | 6D5 | 115546 – Biolegend |
| IgM | BV786 | II/41 | 743328 - BD OptiBuild |
| IgD | BV650 | 11-26c.2a | 405721 – Biolegend |

**Cell proliferation marker**

| Antibody | Fluorophore | Clone | Catalog# & source |
|---|---|---|---|
| Ki67 | FITC | 11F6 | 151211 – Biolegend |

***in vitro***

| Antibody | Fluorophore | Clone | Catalog# & source |
|---|---|---|---|
| B220 | FITC | RA3-6B2 | 103206 – Biolegend |
| CD19 | BV510 | 6D5 | 115546 – Biolegend |
| CD43 | APC-Cy7 | 1B11 | 121220 – Biolegend |
| CD93 | PE-Cy7 | AA4.1 | 136506 – Biolegend |
| CD115 | BV786 | AFS98 | 750888 - BD OptiBuild |
| CD11b | APC | M1/70 | 101212 – Biolegend |
| Gr1 | PE | RB6-8C5 | 108408 – Biolegend |

**Cell Surface Markers used for Characterizing Hematopoietic Cells**

| Cell Population | Surface Markers |
|---|---|
| LT-HSC | $Lin^-$ $c\text{-}Kit^+$ $Sca\text{-}1^+$ $Flk2^-$ $CD48^-$ $CD150^+$ |
| ST-HSC | $Lin^-$ $c\text{-}Kit^+$ $Sca\text{-}1^+$ $Flk2^-$ $CD48^-$ $CD150^-$ |
| MPP2 | $Lin^-$ $c\text{-}Kit^+$ $Sca\text{-}1^+$ $Flk2^-$ $CD48^+$ $CD150^+$ |
| MPP3 | $Lin^-$ $c\text{-}Kit^+$ $Sca\text{-}1^+$ $Flk2^-$ $CD48^+$ $CD150^-$ |
| MPP4 | $Lin^-$ $c\text{-}Kit^+$ $Sca\text{-}1^+$ $Flk2^+$ $CD48^+$ $CD150^-$ |
| CLP | $Lin^-$ $IL7R^+$ $c\text{-}Kit^+$ $Sca\text{-}1^+$ |
| ALP | $Lin^-$ $IL7R^+$ $c\text{-}Kit^+$ $Sca\text{-}1^+$ $Ly6D^-$ |
| BLP | $Lin^-$ $IL7R^+$ $c\text{-}Kit^+$ $Sca\text{-}1^+$ $Ly6D^+$ |
| CMP | $Lin^-$ $c\text{-}Kit^+$ $Sca\text{-}1^-$ $CD16/CD32^-$ $CD34^+$ |
| GMP | $Lin^-$ $c\text{-}Kit^+$ $Sca\text{-}1^-$ $CD16/CD32^+$ $CD34^+$ |
| MEP | $Lin^-$ $c\text{-}Kit^+$ $Sca\text{-}1^-$ $CD16/CD32^-$ $CD34^-$ |
| Pre CFU-E | $Lin^-$ $c\text{-}Kit^+$ $Sca\text{-}1^-$ $CD41^-$ $CD16/CD32^-$ $CD150^+$ $CD105^+$ |

| | |
|---|---|
| Pre Meg-E | Lin$^-$ c-Kit$^+$ Sca-1$^-$ CD41$^-$ CD16/CD32$^-$ CD150$^+$ CD105$^-$ |
| **Bone marrow B cell** | |
| B cells | CD3$^-$ CD11b$^-$ GR1$^-$ TER119$^-$ B220$^+$ |
| Pre Pro-B | CD3$^-$ CD11b$^-$ GR1$^-$ TER119$^-$ B220$^+$ CD43$^+$ CD19$^-$ CD93$^+$ |
| Pro-B | CD3$^-$ CD11b$^-$ GR1$^-$ TER119$^-$ B220$^+$ CD43$^+$ CD19$^+$ CD24$^{med}$ |
| Early Pre-B | CD3$^-$ CD11b$^-$ GR1$^-$ TER119$^-$ B220$^+$ CD43$^+$ CD19$^+$ CD24$^{high}$ |
| Late Pre-B | CD3$^-$ CD11b$^-$ GR1$^-$ TER119$^-$ B220$^+$ CD43$^-$ IgM$^-$ IgD$^-$ |
| Immature B | CD3$^-$ CD11b$^-$ GR1$^-$ TER119$^-$ B220$^+$ CD43$^-$ IgM$^{med}$ IgD$^-$ |
| Transitional B | CD3$^-$ CD11b$^-$ GR1$^-$ TER119$^-$ B220$^+$ CD43$^-$ IgM$^{high}$ IgD$^-$ |
| **Spleen B cell** | |
| B cells | B220$^+$ |
| Transitional B | B220$^+$ CD19$^+$ CD23$^+$ CD93$^+$, |
| Transitional 1 (T1) | B220$^+$ CD19$^+$ CD23$^{low}$ CD93$^+$ IgM$^{high}$ |
| Transitional 2 (T2) | B220$^+$ CD19$^+$ CD23$^{high}$ CD93$^+$ IgM$^{high}$ |
| Transitional 3 (T3) | B220$^+$ CD19$^+$ CD23$^{high}$ CD93$^+$ IgM$^{low}$ |
| Mature B | B220$^+$ CD19$^+$ CD23$^+$ CD93$^-$ |
| Follicular B | B220$^+$ CD19$^+$ CD23$^+$ CD93$^-$ IgM$^+$ CD21/35$^{low}$ |
| Marginal zone B | B220$^+$ CD19$^+$ CD23$^+$ CD93$^-$ IgM$^+$ CD21/35$^{high}$ |
| B1 cells | B220$^+$ CD19$^+$ CD23$^-$ CD43$^+$ |

| **Lymph nodes and peripheral blood B cells** | |
|---|---|
| B cells | B220$^+$ CD19$^+$ |

***Table S3.*** List of qRT-PCR Primers

| Genes | Probe / primer | Sequence (5’ → 3’) | Source |
|---|---|---|---|
| *Mllt1* | Probe | /56-FAM/CCTAAACCC/ZEN/AAGCGTGTGTGCAAG/3IABkFQ/ | IDT |
| | Primer 1 | CTGAGCAGTGTGACATCCAG | |
| | Primer 2 | CCGACTCCTCCACTTTGTAG | |
| *Polr2a* | Probe | /5HEX/CCACCACCT/ZEN/CTTCCTCCTCTTGC/3IABkFQ/ | IDT |
| | Primer 1 | GGTCCTTCGAATCCGCATC | |
| | Primer 2 | CAGGGTCATATCTGTCAGCATG | |
| *Pax5* | Probe | /56-FAM/AAGCTGCCT /ZEN/GGAGATGTCGCA /3IABkFQ/ | IDT |
| | Primer 1 | CCAGATGTAGTCCGCCAAAG | |
| | Primer 2 | GCTTGATGCTTCCTGTCTCAT | |
| *Ebf1* | Probe | /56-FAM/AGGCGCACA /ZEN/ TAGAAATCCTGTTCTGT/3IABkFQ/ | IDT |
| | Primer 1 | CCAACAGCGAAAAGACCAAT | |
| | Primer 2 | GCTTGTTTTGTCATGGAGTCG | |
| *Tcf3* | Probe | /56-FAM/TCCTGGACT /ZEN/ TCAGCATGATGTTCCC/3IABkFQ/ | IDT |
| | Primer 1 | GCTCTGACAAGGAACTGAGTG | |
| | Primer 2 | AGTCCTGAGCCTGCAAAC | |
| *Foxo1* | Probe | /56-FAM/ATTGAATTC /ZEN/ TTCCAGCCCGCCGA/3IABkFQ/ | IDT |
| | Primer 1 | CTTCAAGGATAAGGGCGACAG | |

| | | | |
|---|---|---|---|
| | Primer 2 | AGTTCCTTCATTCTGCACTCG | |
| *Irf8* | Probe | /56-FAM/ ACCTCCTGA/ZEN/TTGTAATCCTGCTTGCC /3IABkFQ/ | IDT |
| | Primer 1 | GAAGACCATGTTCCGTATCCC | |
| | Primer 2 | TGTCTCCCTCTTTAAACTTCCC | |
| *IL7r* | Probe | /56-FAM/ATCTTCTAG/ZEN/GTCTCCATCCTGGGCA /3IABkFQ/ | IDT |
| | Primer 1 | GCTTTCGCTATAGTTTTCTGCTT | |
| | Primer 2 | TGACTTCCATCCACTTCCAAC | |
| *Blk* | Probe | /56-FAM/AGTTCTTCA/ZEN/ACCACACAGACCCTGG/3IABkFQ/ | IDT |
| | Primer 1 | GCTTCTGGTTGCCGCTA | |
| | Primer 2 | TCTCACTGACCTGCCTCTT | |
| *Bach2* | Probe | /56-FAM/CCCAGTTCC/ZEN/CAGCATCCTAAGACT/3IABkFQ/ | IDT |
| | Primer 1 | TGTAGCCTTCTCATCTCTTCCT | |
| | Primer 2 | ATCCACAGACATGCCGTTC | |
| *Vpreb1* | Probe | /56-FAM/AGAAGATGC/ZEN/TAATGGTGGCTGATGCA/3IABkFQ/ | IDT |
| | Primer 1 | CTCATGCTGCTGGCCTAT | |
| | Primer 2 | TGTTATGGTCGTTGCTCAGG | |
| *Igll1* | Probe | /56-FAM/CCACAGCTC/ZEN/TGCTCCTTTCTGCT/3IABkFQ/ | IDT |
| | Primer 1 | CCACCACATACTTTCCCCAA | |
| | Primer 2 | CGTGGGATGATCTGGAACAG | |

| | | | |
|---|---|---|---|
| *CD24a* | Probe | /56-FAM/TGCTCCTAC/ZEN/CCACGCAGATTTACTG/3IABkFQ/ | IDT |
| | Primer 1 | GACATGGGCAGAGCGATG | |
| | Primer 2 | GGTGCAACAGATGTTTGGTT | |
| *Ikzf1* | Probe | /56-FAM/TCCCAAGTT/ZEN/TCAGGAAAGGAGAGCC/3IABkFQ/ | IDT |
| | Primer 1 | CGCACAAATCCACATAACCTG | |
| | Primer 2 | TCATCTGGAGTGTCACTGACT | |
| *Cebpb* | Probe | /56-FAM/ACACGGGAC/ZEN/TGACGCAACACA/3IABkFQ/ | IDT |
| | Primer 1 | CCGCAGGAACATCTTTAAGTGA | |
| | Primer 2 | GTTTCGGGACTTGATGCAATC | |
| *Runx2* | Probe | /56-FAM/CTACAACCT/ZEN/TGAAGGCCACGGGC/3IABkFQ/ | IDT |
| | Primer 1 | CAACTTCCTGTGCTCCGT | |
| | Primer 2 | CCATCTGGTACCTCTCCGA | |
| *Cebpa* | Probe | /56-FAM/CGCAAGAGC/ZEN/CGAGATAAAGCCAAAC/3IABkFQ/ | IDT |
| | Primer 1 | ACAAGAACAGCAACGAGTACC | |
| | Primer 2 | TCATTGTCACTGGTCAACTCC | |
| *Pu.1* | Probe | /56-FAM/ACGCCTGTA/ZEN/ACATCCAGCTGAGC/3IABkFQ/ | IDT |
| | Primer 1 | TGATCCCCCACCGAAGCA | |
| | Primer 2 | CGTAAGTAACCAAGTCATCCGA | |
| *Gata2* | Probe | /56-FAM/TTCTGGCAG/ZEN/CAGACAGCCTCC/3IABkFQ/ | IDT |
| | Primer 1 | TCGTCTGACAATTTGCACAAC | |
| | Primer 2 | ACAAGATGAATGGACAGAACCG | |

| | | | |
|---|---|---|---|
| *Tcf4* | Probe | /56-FAM/CCAGTGAAA/ZEN/TGTCCACTCGCCAAAG/3IABkFQ/ | IDT |
| | Primer 1 | GATGTTTTCGCCTCCTGTAAG | |
| | Primer 2 | ACCCTGAGCTACTTCTGTCTT | |
| *Itgam* | Probe | /56-FAM/TCTTCTGGT/ZEN/CACAGCCCTAGCCT/3IABkFQ/ | IDT |
| | Primer 1 | CCACAGTTCACACTTCTTTCAG | |
| | Primer 2 | TGTCCAGATTGAAGCCATGA | |
| *Csf1r* | Probe | /56-FAM/CACAGTTTG/ZEN/GCATGGTCAGGGG/3IABkFQ/ | IDT |
| | Primer 1 | CTGGAGGCTATGGAGTTGG | |
| | Primer 2 | CACTAGGCTCGATGACAGG | |
| *Baffr* | Probe | /56-FAM/AGGCTGCTT/ZEN/GTATGTCCAGTGTCC/3IABkFQ/ | IDT |
| | Primer 1 | TGTCCTGTGAGCTCTTCCA | |
| | Primer 2 | CCAGTATCAGTCCCAGGAGT | |
| *S1pr1* | Probe | /56-FAM/AGAAACAGC/ZEN/AGCCTCGCTCAAGC /3IABkFQ/ | IDT |
| | Primer 1 | GGTGTAGACCCAGAGTCCT | |
| | Primer 2 | GTCAGCGAGCAATCCAATG | |

***Table S4.*** List of Primers used for Introducing Mutations in MLLT1 by Site-Directed Mutagenesis

| **Inserted mutations** | **Forward and Reverse primers** |
|---|---|
| MLLT1 Y78A<br>Ac/Cr binding site mutation | Fwd: 5'-<br>catgatgaagccagcggcccccgactcctctact-3'<br>Rev: 5'-<br>agtagaggagtcgggggccgctggcttcatcatg-3' |
| Primer combo 1: MLLT1 R63E<br>K67E<br>RNA binding site mutations | Fwd: 5'-<br>agggggggctcctcgcacacgcgctcgggcttggggaa-3'<br>Rev: 5'-<br>ttccccaagcccgagcgcgtgtgcgaggagcccccct-3' |
| Primer combo 2: MLLT1 R63E<br>K67E<br>RNA binding site mutations | Fwd: 5'-<br>agggaggctcctcgcacacgcgctcgggcttagggaa-3'<br>Rev: 5'-<br>ttccctaagcccgagcgcgtgtgcgaggagcctccct-3' |
| MLLT1 D536R<br>AF4 binding site mutation | Fwd: 5'-<br>ggagaagaggtcgaagcggaaggtggtgttggtg-3'<br>Rev: 5'-caccaacaccaccttccgcttcgacctcttctcc-3' |
| MLLT1 D538R<br>DOT1L binding site mutation | Fwd: 5'-<br>gtccagggagaagaggcggaagtcgaaggtggtg-3'<br>Rev: 5'-caccaccttcgacttccgcctcttctccctggac-3' |

| | |
|---|---|
| MLLT1 E523R<br><br>BCOR binding site mutation | Fwd:<br><br>5'attgaagtggccagtctccctgatcagattcacaatctgc-<br><br>3'<br><br>Rev: 5'-<br><br>gcagattgtgaatctgatcagggagactggccacttcaat-3' |

*Supplementary Methods*

#### Mice

*Mllt1* conditional knock-out mice were generated by introducing *Mllt1* targeting vector (created by International Mouse Knockout Project (KOMP) (PG00238_Z_8_F03)) into ES cells and the resulting ES cells with insert were introduced into blastocysts to give rise to chimeric animals with the targeted *Mllt1* allele. Offspring were bred with animals expressing the Flp recombinase to excise a lacZ-neo cassette yielding mice heterozygous for an *Mllt1* allele with a floxed third exon. Backcrosses were performed between $Mllt1^{fl/fl}$ homozygous mice and hemizygous $Rosa26Cre\text{-}ER^{T2}$ animals to produce $Mllt1^{fl/fl}$*;* $Rosa26Cre\text{-}ER^{T2/+}$ mice (experimental) or $Mllt1^{fl/fl}$*;* +/+ (control). Genotyping primers are listed in *Online Supplementary Table S1*. Administration of tamoxifen to adult animals result in cre-mediated excision of the *Mllt1* third exon and absence of *Mllt1* transcripts in experimental mice. Intraperitoneal tamoxifen injections in $Mllt1^{fl/fl}$*;* $Rosa26Cre\text{-}ER^{T2/+}$ mice and $Mllt1^{fl/fl}$ control mice (7-9 weeks old) were 100 mg/kg for 2 consecutive days in a week for 2 weeks. Multiple timepoints, starting at two weeks post tamoxifen, were used for immunophenotype analysis. All animal studies were performed according to standards set forth in the NIH guidelines and approved by Loyola University's Institutional Animal Care and Use Committee.

#### Flow cytometry

For *in vivo* experiments, bone marrow, spleen, lymph nodes and peripheral blood of the mice were harvested and processed. Isolated cells were incubated with 1X RBC lysis buffer (0.155M $NH_4Cl$ + 0.012M $KHCO_3$ + 0.00013M EDTA) for 10 min and 1:500 dilution of Live/Dead Zombie Violet dye for 30 min to eliminate red blood cells and dead cells respectively. Cells were then washed with FACS buffer (PBS+2%FBS) and incubated with fluorescent conjugated-lineage-specific antibodies (*Online Supplementary Table S2*) in FACS buffer at 4C for 30 min. After incubation, cells were washed and followed with subsequent additional steps of mitotracker and/or proliferation marker staining, as appropriate. For *in vitro* experiments, cultured cells were stained for Live/Dead Zombie Violet dye (Biolegend, cat# 423114) followed by incubation with appropriate fluorescent-conjugated antibodies in brilliant stain buffer (BD Horizon, cat# 563794) or FACS buffer at 4C. The incubated *in vivo* and *in vitro* cells were finally analyzed by flow cytometry. All flow cytometry data were acquired using LSR Fortessa cytometer (Becton Dickinson) followed by analysis using FlowJo v9 and v10 software. Cell sorting was performed by flow cytometry core staff using Cytek Aurora CS.

#### Cell proliferation assay

Cell proliferation was assessed using Intracellular Ki67 staining. Following the sequential incubation steps with viability dye, fluorescent labeled antibodies and fixation/permeabilization step (BD Cytofix/Cytoperm kit, cat# 554714), cells were incubated with FITC-Ki67 (Biolegend, cat# 151211; clone: 11F6) at the final step before proceeding flow cytometry data acquisition.

#### Mitotracker staining

After staining bone marrow cells with fluorescent conjugated antibodies specific for lineage markers, cells were washed in PBS and incubated in MitoTracker staining solution containing 75nM MitoTracker Green (Fisher; #M7514), 50nM MitoTracker Red CMXRos (Fisher; M7512), and 50μM verapamil (Sigma; #V4629) in FACS buffer at 37C for 30min. Mitochondrial mass and mitochondrial membrane potential were assessed using the median fluorescence intensity of MitoTracker Green and MitoTracker Red CMXRos, respectively.

##### *in vitro* progenitor B cell differentiation assay

Bone marrow cells from *Mllt1$^{fl/fl}$; Rosa26Cre-ER$^{T2/+}$* mice or *Mllt1$^{fl/+}$; Rosa26Cre-ER$^{T2/+}$* were enriched for B-enriched lineage negative cells by negative selection using MagniSort mouse B cell enrichment kit (ThermoFisher scientific, cat# 8804-6827-74) as per manufacturer's protocol. The biotinylated antibody cocktail in the kit eliminates T cells, monocyte and macrophages, NK cells, granulocytes and erythrocytes. 100,000-500,000 of the resulting B-enriched cells were equally plated between control (400nM ethyl alcohol) and knock-out (400nM 4-hydroxy tamoxifen (4OHT)) conditions in 24-well plate in the IMDM media (Corning, cat# 15-016-CV) supplemented with 10% FBS, $5*10^{-5}$ M of 2-mercaptoethanol, 1mM glutamine, 0.03% peptone primatone (Sigma-Aldrich, cat# P4963), 1% PenStrep (P/S), 10ng/mL IL-7 (PeproTech, cat# 217-17), 100 ng/mL SCF (PeproTech, cat# 250-03) and 50 ng/mL Flt3L (PeproTech, cat# 250-31).

##### RNA sequencing

For *in vitro* cultured cells, total RNA was extracted from each of vehicle- and 4OH tamoxifen-treated B-enriched lineage negative bone marrow cells of three independent biological samples using TRI Reagent (Sigma-Aldrich, #T9424), according to the manufacturer's instructions. Similarly for *in vivo* samples, RNA was isolated from FACS isolated mouse bone marrow-derived B progenitors and CLP from each of three tamoxifen treated *Mllt1$^{fl/fl}$; Rosa26Cre-ER$^{T2/+}$* and *Mllt1$^{fl/fl}$* mice. RNA sequencing for *in vitro* samples was performed by Novogene (Sacramento, CA), and for *in vivo* samples were prepared by the University of Miami Hussman Institute for Human Genomics Sequencing Core.

RNA concentration and purity were assessed with the Agilent 5400, and RNA integrity was confirmed using a Bioanalyzer 2100 system (Agilent Technologies, CA). Samples with RNA Integrity Numbers (RIN) > 8 were used for RNA sequencing. mRNA was isolated from total RNA and used for cDNA library preparation.

For *in vitro* cultured cells, libraries were sequenced using Illumina NovaSeq 6000 generating paired-end 150 bp reads with an average depth of 40 million reads per sample. Raw reads were quality-checked and trimmed using fastp, then aligned to the mm39 mouse genome using HISAT2 v2.0.5 [1]. Gene expression levels were quantified with featureCounts v1.5.0-p3, and differential expression analysis was performed using DESeq2 R package (1.20.0). Genes with an adjusted p-value ≤ 0.05 were considered significantly differentially expressed. KEGG pathway enrichment analyses were conducted using ShinyGO v.0.85 [2] and clusterProfiler R package. Gene set enrichment analysis (GSEA) was performed using the Broad Institute's GSEA 4.3.3 software [3].

For in vivo samples, Second-stranded libraries with ERCC spike-in controls (ThermoFisher Scientific, #4456740) were prepared by the University of Miami Hussman Institute for Human Genomics Sequencing Core using the Nugen Ovation SoLo RNA-seq kit with 15-18 cycles of PCR. Libraries were sequenced to a depth of >30M reads on the NovaSeq X Plus with 150 bp paired-end sequencing.

**RNA-seq alignment:**

Using Cutadapt (v2.6), adapters were removed from both reads [4]. Reads were aligned to the mm10 reference genome using the STAR aligner (v2.7.3), specifying the following parameters: STAR --runThreadN 5 --genomeDir genome_path --readFilesIn trim_R1_fullpath trim_R2_fullpath --outFileNamePrefix name_ --outFilterType BySJout --outFilterMultimapNmax 20 --alignSJoverhangMin 8 --alignSJDBoverhangMin 1 --outFilterMismatchNmax 999 --alignIntronMin 20 --alignIntronMax 1000000 --alignMatesGapMax 1000000 --readFilesCommand gunzip -c --outSAMtype BAM SortedByCoordinate --outWigType bedGraph --outWigStrand Stranded --outWigNorm RPM --alignEndsType EndToEnd [5].

**Differential gene-expression analysis:**

Gene counts were calculated using featureCounts (v2.0.6) [6] with the -p --countReadPairs -s 1 options and the mm10 Gencode annotation file (vM25) that also contained the ERCC spike-ins. Differential gene expression analysis was performed using the package DESeq2 (v1.30.0) with the DESeqDataSetFromHTSeqCount and DESeq functions and data of RNA isolation as a cofactor in the design [7]. The log-fold change was corrected using the ashr method [8]. Significant genes were defined as having a fold change >1.5 and p-adjusted <0.05. Regularized log-counts (rld) and the Wald-statistic were generated with DESeq2. For GSEA (v4.4.0) the Wald statistic ranked list was used with the Mouse_Gene_Symbol_Remapping-Human_Orthologs_MSigDB.v7.4 chip and the c2.all.2026 curated gene set using the weighted enrichment score [9].

**Quantitative RT-PCR**

Total RNA was isolated using TRI Reagent (Sigma-Aldrich, #T9424) and reverse transcription was performed using a High-Capacity cDNA Reverse Transcription Kit (Applied Biosystems, #4368813, Carlsbad, CA). Quantitative RT-PCR (qRT-PCR) was performed using TaqMan probes for the genes listed in *Online Supplementary Table S3* (IDT, Coralville, IA) and TaqMan Fast Advanced Master Mix (Applied Biosystems, #4444557). mRNA expression was normalized to *Pol2ra*, and data were analyzed using the $2^{-\Delta\Delta Ct}$ method [10].

**Seahorse Assay**

Seahorse XF96 plates (Agilent; #101085-004) were prepared with Cell-Tak coating (Corning; #354240) and stored at 4C until the time of the assay. Lineage negative cells were isolated from the bone marrow of *Mllt1*$^{del}$ and control mice using the MojoSort Mouse Hematopoietic Progenitor Cell Isolation Kit (BioLegend; #480004) following the manufacturer's protocol. The lineage negative cells were then resuspended in XF RPMI supplemented with 10mM glucose, 1mM pyruvate, and 2mM L-glutamine (Agilent;

#103681-100). Prior to the assay five replicates of 50,000-75,000 lineage negative cells from *Mllt1*$^{del}$ and control were plated and incubated at 37C in the absence of $CO_2$ for 45min. Cellular oxygen consumption rate (OCR) was measured using the Seahorse XFe96 Analyzer (Agilent) at baseline and after the addition of Oligomycin [1.5μM], FCCP [2μM], and Rotenone/Antimycin A [0.5μM].

**Molecular cloning of MLLT1 and its mutants**

FLAG-tagged MLLT1 and its mutants Y78A and R63E K67E in pTwist-CMV puro vector (Twist Bioscience, cat# 103311) were subjected to restriction digestion using XhoI and EcoRI enzymes which was then gel purified to give the size of ~1.67 kb band of MLLT1 (1-559 aa) along with FLAG-tag. Similarly, MSCV-Cherry (mCherry) vector was linearized using XhoI and EcoRI enzymes and treated with Calf-intestinal alkaline phosphatase (CIP) (New England Biolabs, cat# 0591201) in the concentration of 0.5U/μg of DNA at 37C for 10 min followed by heat inactivation at 80C for 2 min to remove 5'-phosphate groups from DNA. Then, mCherry was ligated with MLLT1 to give the ~ 7.9 kb of vector with MLLT1 followed by transformation of DH5α bacteria with this construct, plasmid preparation using QIAGEN plasmid midi kit (QIAGEN, cat# 12143) and sequenced (ACGT, Inc.) to verify the insertion and mutations. Following the confirmation of insertion and mutations, retrovirus carrying these plasmids were generated and appropriate titrations were made and analyzed by flow cytometry to choose the high titer retrovirus for downstream application [11].

***in vitro* site-directed mutagenesis**

The mutations in MLLT1 (Y78A, R63E K67E, D536R, D538R, E523R) were performed by following the manufacturer's protocol of The QuikChange Lightning Site-Directed Mutagenesis Kit (Agilent Technologies, cat# 210518). In short, the primers for *in vitro* site-directed mutagenesis of each mutation were designed using their primer design guidelines and web-based QuikChange Primer Design Program (*Online Supplementary Table S4*). Once the primers for introducing mutations were synthesized, the mutagenesis was performed using the suggested optimized cycling parameters followed by digestion with Dpn I and transformation of tetracycline and chloramphenicol resistant XL10-Gold ultracompetent bacterial cells (*E. coli* strain engineered by Agilent). Later, plasmid was purified and sequenced, as described above, for each mutation.

***in vitro* complementation assay**

B-enriched lineage negative bone marrow cells were collected and processed as previously explained. B-enriched cells were transduced with retroviruses carrying the following vectors: MLLT1, MLLT1-Y78A, MLLT1-R63E K67E, MLLT1-D536R, MLLT1-D538R and MLLT1-E523R. The vector used for introducing MLLT1 is MSCV-mCherry. For *in vitro* MLLT1 and its mutants' complementation assay, B-enriched cells were transduced using the retrovirus carrying each of the above mentioned MLLT1 and its mutants' vector by spinoculation for 3 hours at 33C and sorted for mCherry-positive cells at 72 hours post transduction and cultured in equal numbers for 400nM ethyl alcohol (control) and 400nM 4-hydroxy tamoxifen (Mllt1 deletion) treatment for two weeks. After two weeks, cells were analyzed in flow cytometry.

#### Immunoglobulin isotyping

Mouse immunoglobulin isotyping was performed using LEGENDplex (6-plex) bead-based assay (Biolegend, cat# 740493) following manufacturer's protocol. Mechanistically, biotinylated beads in this assay kit are specific for capturing IgG1, IgG2a, IgG2b, IgG3, IgA, IgM that are varied by size and internal fluorescence intensities which aids the immunotyping by flowcytometry. Plasma from control and *Mllt1$^{del}$* mice were collected post tamoxifen treatment and stored at -80C. As per manufacturer's instructions, samples and standards were subjected to 50,000 fold dilution and incubated with capture beads and assay buffer in plate shaker followed by a wash with 1X washing buffer and incubation with biotinylated detection antibodies in plate shaker. Following the final wash step, samples were run in flow cytometry and analyzed using Biolegend's LEGENDplex data analysis software (https://legendplex.qognit.com).
